# Evaluating the effects of CD8/CD4 on T cell function in terms of TCR–pMHC–coreceptor catch and slip bonds

**DOI:** 10.64898/2025.12.01.691627

**Authors:** Stefano Travaglino, Amir Hossein Kazemipour Ashkezari, Menglan Li, Valencia E Watson, Peiwen Cong, Larissa Doudy, Hyun-Kyu Choi, Cheng Zhu

**Affiliations:** Wallace H. Coulter Department of Biomedical Engineering, Georgia Institute of Technology and Emory University, Atlanta, Georgia 30332, USA; Parker H. Petit Institute for Bioengineering and Biosciences, Georgia Institute of Technology, Atlanta, Georgia 30332, USA; George W. Woodruff School of Mechanical Engineering, Georgia Institute of Technology, Atlanta, Georgia 30332, USA; Department of Biochemistry, College of Life Science and Biotechnology, Yonsei University, Seoul 03722, South Korea

**Keywords:** T cell receptor, peptide major histocompatibility complex, co-receptor, CD4, CD8, mechanotransduction, catch bond, slip bond

## Abstract

**Background:** T cells interact with peptide-major histocompatibility complex (pMHC) via the T cell receptor (TCR) and coreceptor CD4 or CD8 depending on the MHC class. These interactions form catch and slip bonds depending on the pMHC activity. Coreceptors and bond profiles impact TCR triggering and antigen discrimination.

**Methods:** Built upon our recent correlative analysis of TCR–pMHC catch bond with T cell function, we analyzed 26 pairs of T-cell–pMHC interactions to compare the correlations of their biophysical metrics with antigen-induced T cell responses in two situations: when the coreceptor is prevented vs permitted to bind pMHC.

**Results:** We found that the force-based metrics of TCR bond with pMHC perform better than parameters measured in the absence of force either in situ at the T cell membrane or in fluid phase using purified ectodomain proteins as predictors of T cell activation and thymocyte selection in both cases when the contributions of coreceptors are absent and present. Moreover, CD8 or CD4 co-engagement with pMHC systematically increases these metrics and increases TCR sensitivity and specificity, indicating coreceptor-mediated amplification of, or conversion to, catch-bonds that enhances mechanical tuning of TCR responses.

**Conclusion:** Our findings highlight the importance of force in antigen recognition by the TCR and reveal that parameters derived from the bond profile, especially in the presence of coreceptor, are more informative predictors of T cell activation compared to conventional affinity-based measurements. These results offer mechanistic insights into the roles of catch bonds and coreceptors in TCR antigen recognition.

## 1 Introduction

The vertebrate animals’ adaptive immune system operates through interactions between receptors and ligands expressed on immune and tissue cells. In a typical human individual, tens of millions of T lymphocyte clones expressing unique T cell antigen receptor (TCR) sequences scan different antigen peptides presented by major histocompatibility complex (pMHC) molecules, endowing the TCR repertoire the ability to recognize various pathogens and cancerous cells (1-4). Antigen recognition is a critical event that leads to activation, proliferation, effector function, differentiation, and memory formation of T cells (5-7). The sensitivity and specificity of antigen recognition can be greatly enhanced by CD4 and CD8, which also bind pMHC class II and I, respectively, but do not interact with the antigen peptide itself (8). Both serving as coreceptors for the TCR, CD4 and CD8 share some similarities (*e*.*g*., binding to MHC (9) and Lck (10-12)), but also possess many quantitative (*e*.*g*., displaying differential occupancy (13) and interaction dynamics (14) with Lck) and qualitative (*e*.*g*., having dissimilar structures (9)) differences. The CD4^+^ and CD8^+^ T cell lineages have distinct functions and roles in the immune system (15, 16). CD4^+^ T cells act as helpers or regulators by producing cytokines and regulating other immune cells. CD8^+^ T cells, by comparison, principally function as killers by delivering cytotoxic enzymes to eliminate infected or abnormal cells.

*In situ* measurements at the T cell surface have demonstrated that CD4 and CD8 can bind to TCR-bound pMHC-II and pMHC-I, respectively, with substantially higher binding affinities (17-21) and longer bond lifetimes (20-22) than those to unbound pMHC-II and pMHC-I, to form cooperative TCR–pMHC-II–CD4 and TCR–pMHC-I–CD8 trimolecular bonds (17-22). These (17-22) and other (14, 23-27) studies suggest two mechanisms of how the coreceptor amplifies the sensitivity and specificity of antigen recognition by the TCR: 1) enhance signaling by bringing Lck to the TCR vicinity, which facilitates phosphorylation of the immunoreceptor tyrosine activation motifs (ITAM) in the CD3 subunits associated with the ligand-binding TCRαβ subunits (28, 29), and 2) strengthen adhesion (23, 30) by forming more bonds and prolonging the time of engagement between TCR and pMHC (17-22, 31). Benefits to T cells from these mechanisms include helping trigger the TCR by surpassing the kinetic proofreading threshold for T cell activation (32-34), increasing TCR sensitivity (19, 20), and amplifying antigen discrimination by the TCR (17, 18, 21, 22). The demonstration of the second mechanism and its benefits was made by using *in situ*, or two-dimensional (2D) kinetics measurements (35, 36) of both TCR–pMHC-II/I and pMHC-II/I–CD4/8 bimolecular interactions as well as TCR–pMHC-II/I–CD4/8 trimolecular interactions across the junctional gap between the membranes of a T cell and a surrogate antigen presenting cell (APC) (17-22). Such 2D kinetic analyses reveal that not only does the synergistic TCR-coreceptor cooperation quantitatively increase the number of bonds and prolong the bond lifetime, but the TCR–pMHC-II/I–CD4/8 trimolecular interactions also change the type of bonds qualitatively. Specifically, the cooperation between TCR and CD8 on thymocyte membrane to bind the same pMHC on the APC was found to convert two bimolecular TCR– pMHC-I and pMHC-I–CD8 slip bonds to a TCR–pMHC-I–CD8 trimolecular catch bond for negative selection ligands but not for positive selection ligands (22), a phenomenon called dynamic catch (22, 37). Slip bonds represent the ordinary response of receptor–ligand bonds to force, the lifetime of which shortens with increasing force (38, 39). By contrast, catch bonds are counter-intuitive in that their lifetime is prolonged by increasing force until excessive force overpowers the bond, reverting it into a slip bond (39-41). While catch bonds have been observed in a wide variety of molecular systems underlying very different biological processes (41), for TCR–pMHC-II/I interactions, catch bonds have been suggested as a biophysical characteristic underlying the ability of force to induce or amplify T cell signaling (22, 42-49). This suggestion has been supported by a thorough correlative analysis between a variety of T cell functions and 55 datasets of bond lifetime *vs* force curves of TCR–pMHC-II/I interactions, including both catch and slip bonds (49, 50).

Building on our recent biophysical models of TCR–pMHC bimolecular catch and slip bonds and their correlation with T cell function (50), here we investigated how catch and slip bonds impact T cell activation and function in the presence of coreceptors by extending the model and applying it to compare TCR interactions with pMHC under conditions where coreceptor co-engagement was either prevented or permitted (Fig. 1A). This was done by using machine learning approaches to evaluate 5 metrics of all 26 pairs of TCR bond profiles without and with coreceptor cooperation and their correlation with biological activity. We also investigated the role that coreceptor plays by ligating with TCR-prebound pMHC to prolong TCR–pMHC engagement time through a tripartite structure and/or to convert the TCR–pMHC bimolecular slip bond to a TCR–pMHC– coreceptor trimolecular catch bond on TCR mechanotransduction and T cell signaling. This was done by performing correlative analyses of T cell function with 9-11 metrics of TCR bonds without and with coreceptor cooperation from both two-dimensional (2D) and three dimensional (3D) measurements in the absence and presence of force. Our findings suggest that force-based metrics serve as better predictors of T cell activation and function in both cases in which the coreceptor cooperation is prevented (50) and permitted (this study), which also reveals mechanistic insights into the roles of catch bond and coreceptor in TCR antigen recognition.

**Figure 1.**
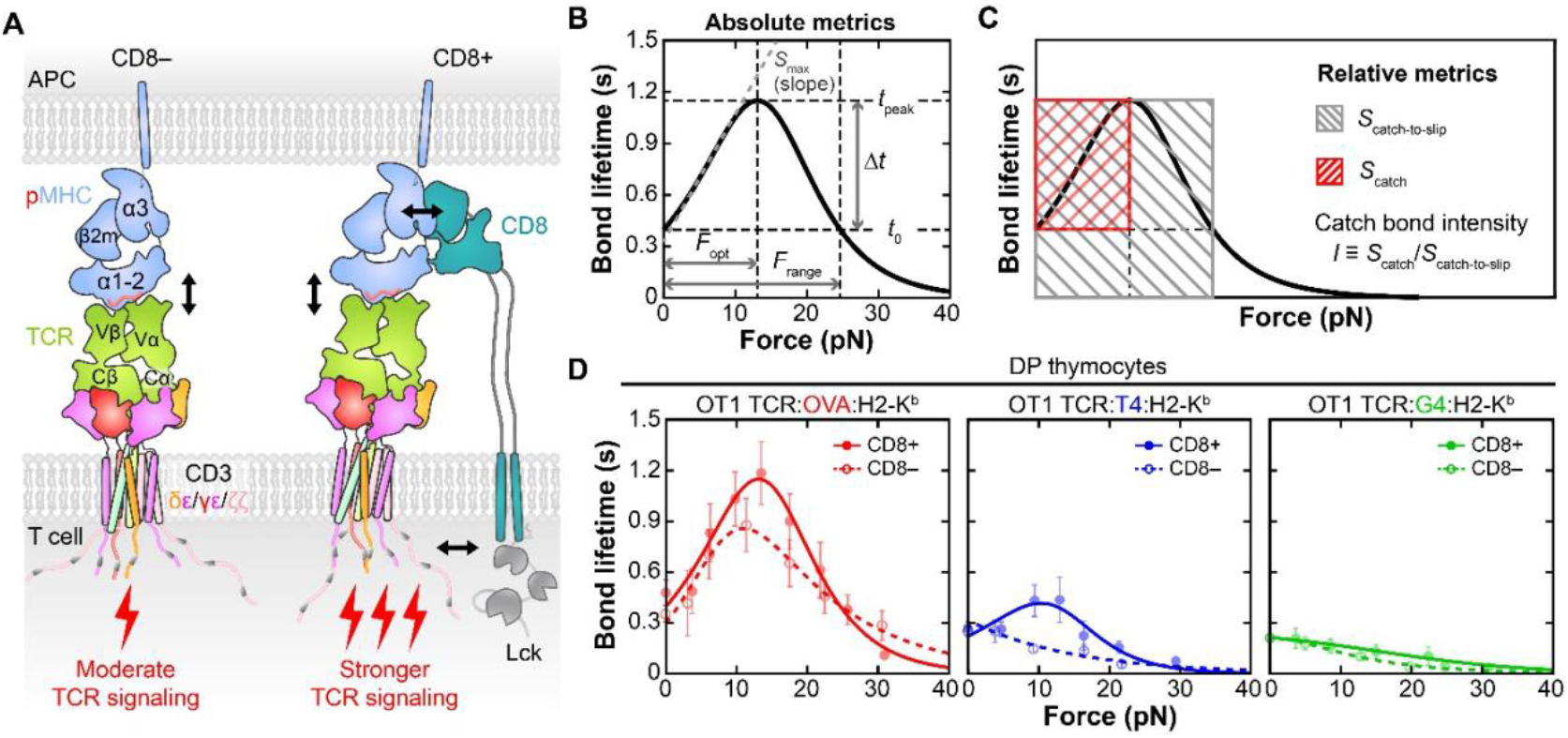
Characterization of TCR catch and slip bonds in the absence and presence of CD8. **(A)** Structural models of αβTCR (*green*)-CD3 (*red, purple*, and *orange* for the indicated subunits) on T cell membrane complexed with a peptide (*light red*) bound MHC-I (*blue*) on APC membrane in the absence (*left*) and presence (*right*) of CD8. Interactions of the pMHC-I with TCR (*both left and right*) and CD8 (*right only*) are indicated by black arrows and the resulting TCR signaling events are indicated by the red lightning signs. Lck, the kinase likely responsible for CD3 phosphorylation is also indicated. (**B)** After smoothing the experimental bond lifetime *vs* force data by fitting it to 1/*k*(*F*) from the conventional model (Methods), five biophysical metrics are calculated for each curve: *t*_peak_ is the peak bond lifetime, Δ*t* is the lifetime increase from the zero-force value *t*_0_ to *t*_peak_, *F*_opt_ is the “optimal force” where lifetime reaches *t*_peak_, and *F*_range_ is the range over which force amplifies lifetime beyond *t*_0_, A*UC* is the area under the bond lifetime *vs* force curve, and *S*_max_ is the maximum slope along the curve. (**C**) Definition of the catch bond intensity *I* as the ratio of two areas from catch-phase (*S*_catch_) and entire-phase (*S*_catch−to−slip_) in force-lifetime profile. (**D**) Fitting of theoretical 1/*k*(*F*) curves to experimental bond lifetime *vs* force data (*points*, Mean ± SEM from *n* > 30 bond lifetime data per force bin, re-analyzed from (22)) of OT1 TCR expressed on CD4^+^CD8^+^ thymocyte T cells interacting with OVA (agonist, *left*), T4 (weak agonist, *middle*) and G4 (antagonist, *right*) peptides presented by WT H-2K^b^ (CD8^+^, dark colored points and curves) or MT H-2K^b^α3A2 (CD8^-^, light colored points and curves).

## 2 Materials and Methods

### 2.1 Isolation of CD8^+^ T Cells from Spleen of OT1 transgenic mice

OT1 transgenic mice (C57BL/6-Tg(TcraTcrb)1100Mjb/J) were sourced from Charles River Laboratory (Lyon, France) and bred in-house at the Georgia Institute of Technology. Both male and female mice, aged between 14 to 16 weeks, were housed under controlled conditions maintaining temperatures between 20°C and 26°C, humidity levels from 40% to 70%, and a 12-hour light-dark cycle.

To isolate naïve OT1 CD8^+^ T cells, mice were euthanized following a protocol approved by the IACUC of Georgia Institute of Technology. The spleens were carefully extracted and mechanically dissociated to create a single-cell suspension, which was then filtered through a 70 μm nylon cell strainer (BD Falcon) to remove debris. Red blood cells were lysed using a lysis buffer, and the remaining splenocytes were washed with the wash buffer (Mouse Erythrocyte Lysing Kit, #WL2000, R&D Systems). CD8^+^ T cells were purified from the splenocyte mixture using an immunomagnetic negative selection kit (cat#19853, STEMCELL Technologies). The isolated CD8^+^ T cells were then resuspended in the culture media (RPMI + 10 % FBS, 100U/ml penicillin and 100 ug/ml streptomycin, 20 mM HEPES, 1mM Sodium pyruvate) and ready for use in subsequent experiments.

### 2.2 Biomembrane Force Probe Assay

The force-dependent kinetics of pMHC dissociation from the TCR, with or without CD8 co-engagement, were assessed using our previously described Biomembrane Force Probe (BFP) assay (22, 42). Briefly, human red blood cells (RBCs) were isolated from whole blood drawn from healthy volunteers using a protocol approved by Institution Review Broad of Georgia Institute of Technology and biotinylated. Streptavidin-coated glass beads were functionalized by incubation with biotinylated pMHC at sub-saturation concentration. Three OT1 TCR cognate peptides presented on two MHC molecules were produced by the NIH Tetramer facility: the wide-type (WT) peptide OVA_323-339_ (SIINFEKL) and two altered peptides, Q4R7 and Q4H7, which were bound to WT mouse MHC H2-K^b^ or a domain swapping mutant (MT) that replaced the mouse α3 domain with that of the human HLA-A2 (H2-K^b^α3A2) to abolish binding of the mouse CD8 (17, 22, 42, 51). RBC was aspirated by a glass micropipette and pressurized. A glass bead was attached to the RBC apex via streptavidin-biotin coupling to assemble a picoforce transducer with a ∼0.3 pN/nm spring constant to interrogate the OT1 CD8^+^ T cell aspirated by an opposing micropipette. Following a controlled contact period to facilitate bond formation between the T cell and pMHC, the T cell was retracted with a predefined ramping force rate (1000 pN/s). In the force-clamp mode of BFP, once a stable bond was established between pMHC and TCR (with or without co-engagement with CD8), T cell retraction was halted to wait for bond dissociation under a constant force. A range of forces was applied to investigate how different force levels modulated the stability and dissociation kinetics of the interaction. Bond lifetime, i.e., the time from force application to bond rupture, was recorded with a temporal resolution of 1000 fps.

Bond lifetimes obtained from the force-clamp assay were plotted against the applied force to generate lifetime-versus-force curves. These curves were analyzed to identify patterns such as catch bonds, where bond lifetime increases with force, and slip bonds, where bond lifetime decreases with force.

### 2.3 Measuring T cell activation stimulated by TCR or both TCR and CD8

T cell activation was measured by imaging of intracellular calcium and staining of activation markers. Glass coverslips (25×75 mm^2^) were sonicated in 50% ethanol for 15 min and rinsed 6X with di-H_2_O. Following rinsing, the coverslips were etched in 250 ml Piranha solution (2:1 ratio of sulfuric acid to H_2_O_2_) for 30 min. Coverslips were washed 6X with di-H_2_O and 3X with 100% ethanol. The surfaces were then silanized with 3% APTES in ethanol for 1 h, then washed 3X in ethanol and dried under argon stream. After drying, one Ibidi Sticky-Slide VI 0.4 was mounted on each coverslip to create 6 microfluidic channels per coverslip while making sure to remove any bubbles from the adhesive areas. NHS-PEG4-Azide was then diluted in 0.1M NaHCO_3_ to 10 mg/ml and 50 μl of the solution added to each channel and incubated at room temperature for 1 h. The channels were then washed 2X with di-H_2_O and 1X with PBS and then blocked with PBS + 0.2% BSA for 0.5 h. During this incubation, the DBCO-bottom strand and the biotin-top strand (56 pN) of TGT probes (220 nM each in 1 M NaCl) were annealed in a thermocycler by heating to 95 °C for 5 min and gradually cooling (−5 °C/min) to 25 °C. After blocking, the channels were washed 3X with PBS and 50 μl of PBS was left in the channels to prevent drying. The TGT probes were then added (50 μl) to each channel and incubated overnight at room temperature to covalently functionalize the channels with TGT probes via strain-promoted alkyne-azide cycloaddition (SPAAC). The following day, channels were rinsed 3X with PBS, and 50 μl of 20 μg/ml streptavidin solution in PBS were added to each channel and incubated for 1 h at room temperature. After rinsing 3X with PBS, 50 μl of 20 μg/ml different types of biotinylated pMHC in PBS + 2% BSA was added to each channel and incubated for 1 h at room temperature. Concurrently, naïve OT1 CD8^+^ T cells at a density of 1 × 10^6^ cells/ml were incubated with 5 μM calcium indicator dye X-Rhod-1 (ThermoFisher Scientific) in R10 medium for 30 min at 37 °C. Cells were washed twice with R10. The channels were washed 2X with HBSS and 1X with HBSS imaging buffer, then added cells onto the ligand-coated surface. Upon addition, intracellular calcium fluxes were imaged under a Zeiss 780 / Elyra PS.1 Superresolution Microscope (OMC) equipped with a ×20 air objective. Cells were excited with a Xenon lamp at 580/15 nm and emission was acquired at 620/60 nm at two frames per second for 12 min.

For the flow cytometry measurements of activation markers on T cell surface, azide-PMMA beads (PolyAn) were coated with 56 pN TGT probes and biotinylated pMHCs by following the protocol described above. Beads were then incubated with cells at a 1:1 ratio. After 6 hours incubation, cells were harvested and stained with PE-anti-CD69 (clone H1.2F3, BDBiosciences, 553237), APC-anti-CD3 (clone 145-2c11, BDBiosciences, 553066), and PC7-anti-CD25 (clone PC61, BDBiosciences, 561780) antibodies for 1 hour at room temperature. After washing twice with FACS buffer (PBS with 2% FBS), cells were analyzed by flow cytometry (BD FACSAria).

### 2.4 Conventional two-pathway model

A simple conventional two-pathway model is (52).

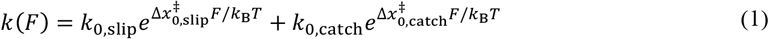

where *F* is applied force, *k*_B_ is Boltzmann constant and *T* is absolute temperature, *k*_0,slip_ and *k*_0,catch_ are the respective zero-force off-rates of the slip- and catch-pathway, 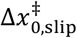 and 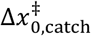 are the respective distances from the bound state to the transition states along the slip- and catch-pathways. Each term follows the Bell equation (38) but the catch pathway parameter 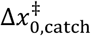 has a negative value to lower off-rate along the path. This is a phenomenological model as it neither considers specific structure and force-induced conformational changes of the interacting molecules nor does it take into account the shifting of the location of the transition state upon force application.

### 2.5 Modified two-pathway model

To allow possible incorporation of specific structure and force-induced conformational changes of the interacting molecules, we modified the above phenomenological two-pathway model (52) by keeping the slip pathway as is but replacing the catch pathway by our recent model for TCR– pMHC-II/I catch bonds (50):

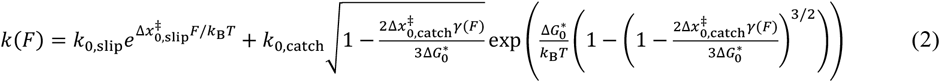

where 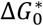 is the differences in free-energy levels along the catch pathway, the structure-based force function *γ*(*F*) scales with the characteristic extension change per unit change of molecular length such that 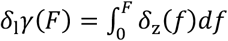. Typically, *δ*_1_ is the partially unfolded extension along the force (*n*^∗^*l*_c_ where *n*^∗^ is the number of amino acids and *l*_c_ = 0.36 nm for unstructured amino acids). Additionally, *δ*_z_(*f*) = *z*(*f*) − *z*_0_(*f*) is the projection on the force direction of the change induced by force *f* of the TCR–pMHC-I/II–CD8/4 extension at the transition state relative to its extension at the bound state (50).

### 2.6 Model fitting of force-dependent bond lifetime profiles of TCR–pMHC-II/I–CD4/8 interactions

Model fitting to experimental data was accomplished through nonlinear curve fitting utilizing the Levenberg-Marquardt algorithm (MATLAB built-in function). In essence, the best-fitting parameter set was obtained by fitting the model to the mean value of bond lifetime *vs* force data. Additionally, the standard error (SE) of fitting was calculated by independently fitting the model to the Mean ± standard error of the Mean (SEM) *vs* force data. It was observed that the parameters fitted to the Mean of bond lifetime were robust and fell within the range of parameters ± SEM. All published experimental bond lifetime *vs* force data were measured at room temperature, as reported in the references (22, 27, 42-44, 48). The residual sum of squares (RSS) and the reduced Chi-squared (*χ*_*ν*_^2^) for comparison of the goodness-of-fit are calculated using their respective definitions: 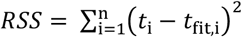 where *t*_i_ is the *i*-th bond lifetime measurement and *t*_fit,i_ is bond lifetime value from fitted curve at each *i*-th force value; 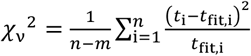 where *n* is number of observations and *m* is the number of parameters used in each model. Any other linear fits shown in figures were performed in MATLAB using the built-in linear regression model. For spline fitting, smoothing splines were applied using moving average filtering, a built-in function in MATLAB.

### 2.7 Defining biophysical metrics from lifetime *vs* force curves

From each best-fit bond lifetime *vs* force curve, we calculated five biophysical characteristics defined in Fig. 1B (50): the peak bond lifetime (*t*_peak_, in seconds), the optimal force where catch-slip bond lifetime peaks ( *F*_opt_, in pN), the catch bond intensity or catchiness ( *I* = Δ*tF*_opt_⁄*F*_range_*t*_peak_, unitless) where Δ*t* is the change from *t*_peak_ to the force-free bond lifetime, *t*_0_ = 1⁄*k*_0_, and *F*_range_ is the force range where bond lifetime returns from *t*_peak_ back to *t*_0_, the total area under curve (*AUC*), and the maximum slope (*S*_max_) along the curve. The dimensionless scaled metric *I* has a geometric meaning as the ratio of two areas: 1) the rectangular area bounded by two horizontal lines *t* = *t*_0_, *t*_peak_ and two vertical lines *f* = 0, *F*_opt_ and 2) the area boxed by *t* = 0, *t*_peak_ and *f* = 0, *F*_range_ (Fig. 1C).

To test whether and, if so, how the above biophysical metrics would be useful in characterizing the TCR–pMHC-II/I–CD4/8 trimolecular interactions, we analyzed a total of 47 data curves published by three labs (22, 27, 42-44, 48) (Supplementary Figs. 2A, 3A, 3B, the blue curve in left panel of 3C, and 3D-3G), plus 5 new data curves generated in this study (Supplementary Fig. 3C except the blue curve in left panel). Datasets for each TCR include one or more pairs of curves, with each pair consisting of a peptide or a panel of peptides presented by either WT or MT MHC allowing or preventing CD8 co-engagement, respectively. Eight MHC class I restricted TCRs totaling 21 pairs of bond lifetime *vs* force curves were analyzed: 10 pairs for the murine OT1 and 2C TCRs expressed on CD4^+^CD8^+^ (double positive, DP) thymocytes interacting with reactive peptides (7 for OT1 and 3 for 2C, Supplementary Fig. 3A and 3B) (22) and 3 pairs for the OT1 TCR expressed on CD8^+^ (single positive, SP) naïve T cells interacting with three of these reactive peptides ( (42) and this study, Supplementary Fig. 3C), which were presented by H-2K^b^ or H-2K^b^α3A2, which were measured by the Zhu lab using the biomembrane force probe (BFP); 7 pairs measured by the Evavold lab using BFP, including the murine P14 TCR expressed on CD8^+^ naïve T cells (48) and four mouse TCRs expressed on hybridomas (27) interacting with their respective specific peptides bound to WT and MT H-2D^b^ (3 for P14 and 1 for each of the other four TCRs, Supplementary Fig. 3D and 3E); and 1 pair measured by the Lang lab with optical tweezers using the murine N15 TCR expressed on CD8^+^ T cell interacting with VSV8 bound to WT and MT H-2K^b^ (Supplementary Fig. 3F) (44). In addition, 2 MHC class II restricted TCRs were analyzed: human E8 TCR interacting with TPI:HLA-DR1 in the absence or presence of CD4 (20) (Supplementary Fig. 2A) and mouse 3.L2 TCR expressing on CD4^-^CD8^+^ or CD4^+^CD8^-^ naïve T cells interacting with 4 peptides presented on I-E^k^ (43, 53)(Supplementary Fig. 3G), all measured by the Zhu lab using BFP.

### 2.8 Data and statistical analysis

Results are presented either as Mean ± SE, Mean ± SD, or as mean only wherever applicable. P values were calculated by either two-sided paired or unpaired t-test with ^*\**^*P* ≤ 0.05, ^**^*P* ≤ 0.01, ^***^*P* ≤ 0.001, ^****^*P* ≤ 0.0001. Uniform Manifold Approximation and Projection (UMAP) analyses were performed in python using the UMAP parameters outlined in figures and figure legends. Custom MATLAB software was written for analysis of BFP data. Statistical details have also been provided in figure legends and Methods where applicable.

## 3 Results

### 3.1 Two-pathway models for catch bonds

Two decades ago, when receptor–ligand catch bonds were first demonstrated (40), several two- and three-pathway kinetic models were developed to provide a theoretical explanation for the biphasic trend of bond lifetime that first increases (catch), and then decreases (slip), with increasing force (52, 54-57). These models are based on the kinetic rate theory and use abstract physical parameters to describe a generic energy landscape (58-61). By comparison, our recent single-pathway models incorporate the structures, elastic properties, and force-induced conformational changes of the TCR–pMHC-I/II complexes at the sub-molecular level, including domain stretching, hinge rotation, and molecular extension (50). A limitation of single-pathway models is the assumption that bonds must dissociate from a single state along a single pathway (54, 62, 63), which is consistent with some but not all TCR catch bond data that we published (22, 42, 43). A piece of evidence for the presence of more than a single bound state and/or a single dissociation pathway comes from the observation that in some cases lifetimes of TCR bonds are distributed as more than a single exponential, which are more apparent for TCR–pMHC-I–CD8 trimolecular interactions than TCR–pMHC-I bimolecular interactions (22) (Supplementary Fig. 1). Two- and three-pathway models overcome this limitation by assigning different subpopulations of bonds associated with distinct exponential lifetime distributions to different dissociation pathways with distinct force-dependent lifetime averages (56, 64, 65). In a recent paper, we used another two-pathway model to globally fit scattered individual bond lifetime *vs* force measurements of TCR– pMHC-II, pMHC-II–CD4, and TCR–pMHC-II–CD4 interactions simultaneously (20).

In this work, we first constructed a new two-pathway model to take advantage of both our recent single-pathway models containing structural information of the TCR–pMHC-II/I molecules (50). This was done by keeping the slip pathway of the conventional two-pathway model (52) as is but replacing the catch pathway by our recent models for TCR–pMHC-II/I catch bonds (50) (Methods). We compared its ability to describe experiments with the simplest conventional two-pathway model capable of accounting for double exponential bond lifetime distributions (52). We found that both models are equally capable of fitting the average bond lifetime *vs* force curves for both TCR–pMHC-II bimolecular and TCR–pMHC-II–CD4 trimolecular interactions (20) and result in nearly identical force-based metrics for catch bonds (Supplementary Fig. 2A and 2B). We further compared fitting of the scattered bond lifetime *vs* force measurements without binning (20), finding that our newly constructed model with structural information did significantly better than the conventional model without structural information by F-test using the *χ*_ν_ ^2^ measure of the goodness-of-fit (Supplementary Fig. 2C and 2D). Compared to the phenomenological model, our structure-based model shows better and similar goodness-of-fit for TCR–pMHC-II interactions in the presence and absence of CD4, respectively, suggesting that trimolecular catch bonds may be better modeled with structural aspects in the model formulation.

### 3.2 Reducing data representation from a force-lifetime curve to biophysical metrics

The main goal of this work was to examine the correlation between T cell function and force-lifetime profile of TCR bond with pMHC in the absence and presence of coreceptor using all published data to date. Except for Rushdi *et al*. (20), however, the literature data are reported as Mean ± SEM of bond lifetime *vs* Mean ± SEM of force, typically consisting of 6-10 force points (Fig. 1D; Supplementary Figs. 2A, 2E, 2F, and 3). This limits our ability to use our new two-pathway model due to the potential problem of over-fitting. Spline smoothing such data generates curves similar to those fitted by the conventional two-pathway model (Supplementary Fig. 2E and F). However, the scattered bond lifetime *vs* force measurements without binning could not be smoothed by spline (Supplementary Fig. 2G and H). Therefore, we used the conventional two-pathway model to fit published data (22, 27, 43, 44, 48), including 21 pairs of TCR–pMHC-I curves with and without CD8 co-ligation (Supplementary Fig. 3A-3F) and 5 pairs of TCR–pMHC-II curves with and without CD4 co-ligation (Supplementary Figs. 2A and 3G), finding excellent agreement in all cases.

After model fitting to smooth the noisy data, we calculated five biophysical characteristics defined in Fig. 1B: *t*_peak_, *F*_opt_, *I, AUC*, and *S*_max_ (50) (Methods). These metrics are calculated to reduce data representation from a force-lifetime curve into a few characteristic values. The reason is, unlike binding affinity which has a single value when measured in the absence of force, catch and slip bonds exhibit varying trends in how lifetime changes with increasing force, and can differ significantly in shape and magnitude. Distilling the continuous curves into discrete features facilitates their comparisons with T cell function.

### 3.3 Evaluating metrics of TCR bond profiles without and with CD8 co-ligation and the correlation with their categorizations of biological activity

We (22, 42, 43, 46) and others (27, 45, 47, 48) have observed that TCR forms pronounced catch bonds with agonist pMHC-I but less pronounced catch bonds and slip bonds with weak agonists and antagonists. When CD8 is allowed to bind pMHC-I together with TCR to permit the possible formation of a synergistic TCR–pMHC-I–CD8 trimolecular bond, the catch bond of the agonist becomes more pronounced, the slip bond of the weak agonist may change to a weak catch bond, and the slip bond of the antagonist may remain as a slip bond (Fig. 1D; Supplementary Fig. 3B and 3C). In our recent correlative analysis of TCR–pMHC bimolecular catch bonds with T cell function, we observed variable degrees of correlation between the biological activity and biophysical metrics (50).

Built upon these studies, we examined the impacts of CD8 on TCR–pMHC bond profiles. We compared 5 biophysical metrics evaluated from 21 pairs of bond profiles of 8 TCRs forming catch or slip bonds with pMHCs without and with CD8 cooperation in Fig. 2A: *F*_opt_ (1^st^ pair), *I* (2^nd^ pair), *AUC* (3^rd^ pair), *t*_peak_ (4^th^ pair), and *S*_max_ (5^th^ pair). By visualizing the heatmap color changes from the CD8- to CD8+ column within each metric and the progressive cooling of colors from red to blue across the five metrics, it is evident that the ability of these metrics to capture features of the bond profiles follow the ranking of *F*_opt_ > *I* > *AUC* > *t*_peak_ > *S*_max_. Compared to limiting T cell interactions with pMHC to TCR only, allowing CD8 to also bind pMHC increased all 5 metrics for all 21 pairs of interactions except for a few cases, as indicated by the color changes from cool to warm for nearly all of the 105 cases (Fig. 2A). Averaging 21 values in each column, CD8 increased the values of the five metrics by 83% for *F*_opt_ (*P* = 0.0006), 73% for *I* (*P* = 0.0056), 80% for *AUC* (*P* = 0.0206), 50% for *t*_peak_ (*P* = 0.0022), and 356% for *S*_max_ (*P* = 0.0693). These quantitative values confirm the qualitative impression from visual inspection of color changes. The results indicate a general trend that permitting CD8 to bind pMHC-I and to form cooperative TCR– pMHC-I–CD8 bonds enlarges the catch regime (increased *F*_opt_), enhances the catchiness of the bond (increased *I*), expands the force range over which bond lifetime is prolonged (increased *AUC*), prolongs lifetime (increased *t*_peak_), and steepens the slope of the bond profile (increased *S*_max_).

**Figure 2.**
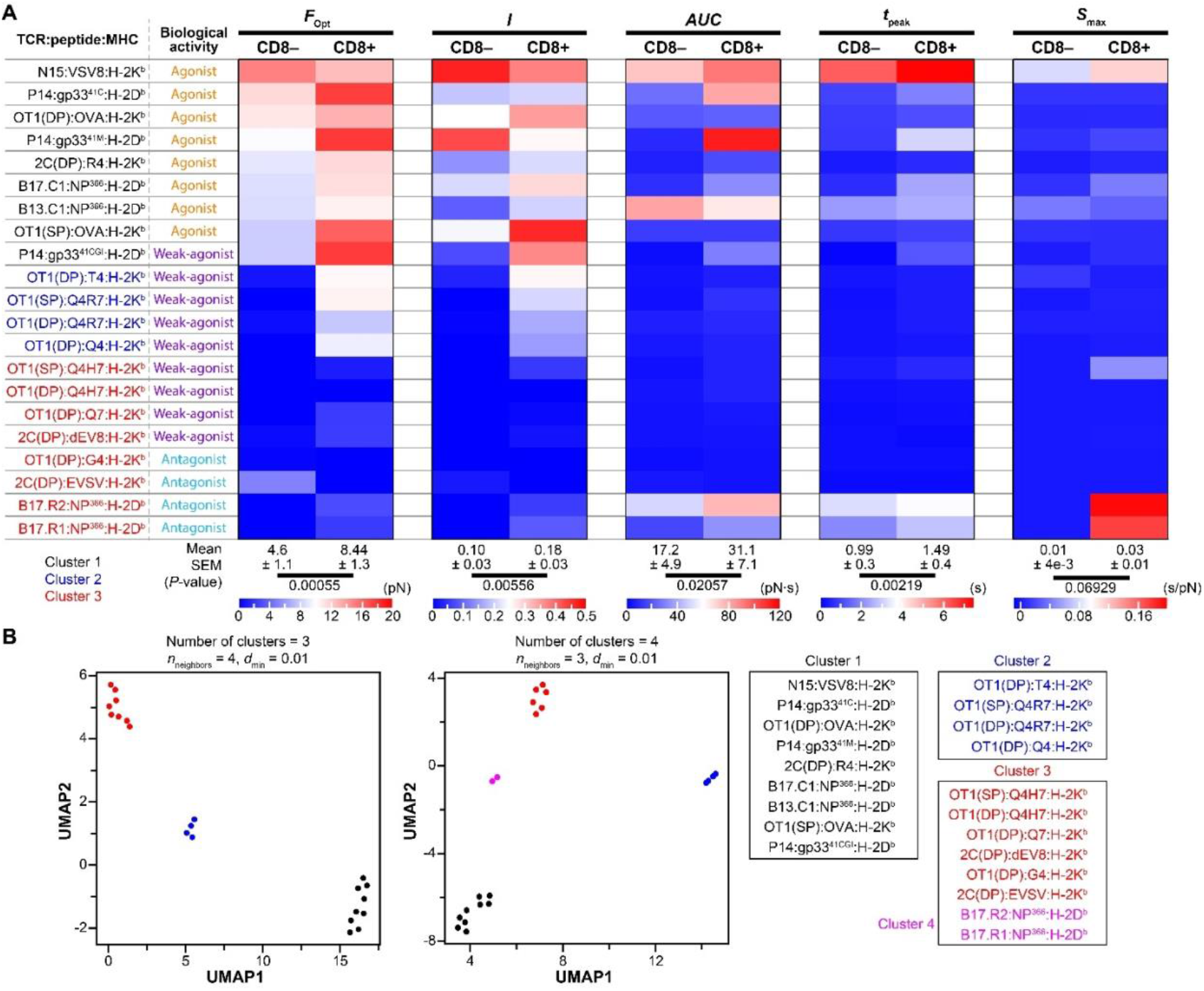
Evaluating metrics of TCR bond profiles without and with CD8 cooperation and their correlation with biological activity. **(A**) *1*^*st*^ *column*: List of 21 pairs of interactions between TCR and pMHC-I, which are grouped into three color coded clusters (cluster 1 = black, cluster 2 = blue, cluster 3 = red), as indicated on the bottom of the 1st row. *2*^*nd*^ *column*: Biological activities of these interactions based on literature, which are grouped into three color-coded categories (agonist = orange, weak agonist = purple, antagonist = cyan). *3*^*rd*^ *– 12*^*th*^ *columns*: Five pairs of metrics (*F*_opt_, *I, AUC, t*_peak_, and *S*_max_, as indicated on the top of the 1st row) of force-dependent lifetime curves of 8 TCRs forming catch and slip bonds with their respective panels of pMHCs without (CD8-) and with (CD8+) coreceptor cooperation. Their values are shown as heatmaps with specific color codes for each metric indicated at the bottom. For each interaction pair, the change in color indicates the change in the metric values between the case in which CD8 was prevented from binding pMHC and the case in which CD8 was permitted to bind pMHC. The mean and SEM values across all 21 pairs of interactions are also shown in the bottom, as well as the P-values that indicate the significance of the increases in the values from the CD8-column to the CD8+ column. (**B**) Uniform Manifold Approximation and Projection (UMAP) analysis was used to group the 21 pairs of interactions into 3 (left) or 4 (right) clusters based on their 10 values from the 5 metrics of each pair of force-lifetime curves measured in the absence and presence of CD8 (shown in the heatmap in A).

It has been proposed that CD8 amplifies TCR signaling and T cell activation, but the degree of amplification varies depending on ligand potency (19, 21, 66, 67). Therefore, we next tested whether this contention was reflected in the above biophysical analysis and, if so, whether it could be recapitulated by our bond profile metrics. To extract the maximum amount information from all 5 (bond profile metrics) × 21 (TCR-pMHC pairs) × 2 (without and with CD8) = 210 metrics values, we performed sequential dimensionality reduction by UMAP and clustering by K-means. We found that the 21 pairs of interactions are well segregated into 3 or 4 clusters within certain parameter ranges (number of neighbors 2 ≤ *n*_*neighbors*_ ≤ 5, minimum distance 0.01 ≤ *d*_min_ ≤ 0.1), as depicted in the UMAP plots (Fig. 2B).

Based on their biological activities reported in the literature, the 21 interactions are categorized as agonists, weak agonists, and antagonists (Fig. 2A). Interestingly, all agonists belong to Cluster 1, all antagonists belong to Cluster 3, and Cluster 2 comprises all weak agonists. Cluster 3 includes as many weak agonists as antagonists, consistent with the imperfect classification between these two categories in the literature (68). The bottom two antagonists become separated from Cluster 3 and form their own Cluster 4 when we decreased the UMAP *n*_*neighbors*_ parameter to 3 (stronger focus on local *vs* global structure). Interestingly, the structures of these two TCR (B17.R1 and B17.R2) complexed with NP^366^:H2-D^b^ display a 180° reverse docking orientation compared to all other interactions analyzed here (27). Cluster 1 also contains the interaction of P14 TCR with the gp33 peptide 11mer 41CGI, which forms a baby catch bond when CD8 was prevented from binding but becomes a bona fide catch bond when CD8 was permitted to bind (48) (Supplementary Fig. 3D, third panel). Functionally, 41CGI is less biologically active than the 9mers 41C and 41M, the latter is created by introducing a single methionine at the carboxy terminal end replacing the cysteine at position 41. Kolawole *et al*. call 41C and 41M as agonist and super agonist, respectively (48), so we label 41CGI as weak agonist to provide a relative grading. On DP thymocytes, 2C TCR formed a slip bond with dEV8 when CD8 was prevented to bind and did not change much when CD8 was allowed to bind (22) (Supplementary Fig. 3A, second panel), but the same interaction showed clear a catch bond when measured using hybridoma cells (46). These results indicate that certain biological features resulted from different interaction pairs are captured by their memberships in the different clusters generated using UMAP analysis of their bond profiles.

### 3.4 Correlating 2D and 3D biophysical metrics measured in the absence and presence of force with biological functions

The UMAP analysis of bond profiles of 21 pairs of interactions groups them into 3 or 4 clusters with distinct functional and/or structural features corresponding to agonists, weak agonists, antagonists, and reversed TCR–pMHC docking orientation (Fig. 2). However, this only maps bond profiles into biological activities categorically but does not decipher the relative contributions from different bond profile metrics to identify the best predictor(s) of T cell response. We next asked which bond profile metric or group of metrics best correlate with T cell response to antigen stimulation when CD8 is either prevented or permitted to cooperate with the TCR. We also asked how the five metrics tested in Fig. 2 compare to other biophysical metrics previously used to correlate T cell activation and function, including four additional 2D kinetic parameters and two 3D biophysical parameters. The first is the average number of bonds, calculated from the steady-state adhesion frequency *P*_a_(∞) = 1 − *e*^−⟨*n*⟩^. per densities of TCR (*m*_TCR_) and pMHC (*m*_pMHC_). When WT MHC was used to permit CD8 binding, the average number of bonds, ⟨*n*⟩ = −ln(1 − *P*_a_), takes into account all bond species, including two TCR–pMHC and pMHC–CD8 bimolecular bonds as well as the TCR–pMHC–CD8 trimolecular bond. After normalizing ⟨*n*⟩ by *m*_TCR_ × *m*_pMHC_, we formulated the first parameter 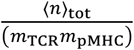, with the subscript “tot” to indicate total bond number. When MT MHC was used to present the peptides and prevent CD8 binding, the average number of bonds contains only the contribution of the TCR–pMHC bimolecular bond, and this parameter is reduced to 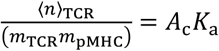, *i*.*e*., the previously defined effective 2D affinity of TCR for the pMHC (35, 36), as indicated by the subscript “TCR”. The second 2D parameter, normalized synergy, is defined as 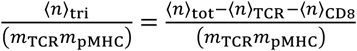, which isolates the contribution from the trimolecular bond, as indicated by the subscript “tri”. Note that normalized synergy vanishes when CD8 is absent. In addition, we calculated two force-based 2D parameters. The first is the average number of total bonds per densities of TCR and pMHC evaluated at *F*_opt_, 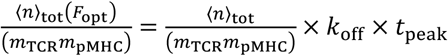, which in the absence of CD8 reduces to effective 2D affinity evaluated at *F*_opt_. This was suggested by our recent finding that *A*_c_*K*_a_(*F*_opt_) best predicted the mechanotransduction of CD40 on B cells (69). The second is the normalized synergy at *F*_opt_, 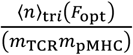, which also vanishes when CD8 is absent. In addition to 2D metrics, we also added two 3D parameters: dwell time *t*_1/2_, and avidity *K*_v_ (reduced to affinity *K*_a_ when CD8 is absent) (29, 70).

In choosing systems for such analyses, we noted some inconsistencies when comparing bond profiles of different TCRs. For example, the two pairs of bond profiles of P14 TCR with 41CGI and OT1 TCR with OVA are very different without CD8 but quite similar with CD8 (compare Supplementary Fig. 3B, first panel with Supplementary Fig. 3D, third panel). Therefore, we chose to focus on a single TCR system, the OT1 TCR interactions with a panel of pMHCs for which both biophysical and biological measurements are available for both cases of with and without CD8. We performed this analysis on data measured using both DP thymocytes and SP naïve T cells. For the former, the biological activities of 7 peptides were measured as EC_50_ from the peptide dose curves of CD69 upregulation (70). For the latter, 4 functional readouts induced by 3 peptides were measured, including the induction of activation markers CD69/CD25/CD3 and intracellular calcium (Supplementary Fig. 4).

We aimed to evaluate the power of various TCR biophysical parameters to predict antigen-induced functions of T-lineage cells under physiological conditions, which occur *in vivo* in the presence, not in the absence, of CD8. Therefore, we plotted three sets of biophysical parameters *vs* function data graphs, respectively from three combinations of measurements made in the presence (+) and absence (-) of CD8: 1) CD8^+^ parameters *vs* CD8^+^ function, 2) CD8^-^ parameters *vs* CD8^+^ function, and 3) CD8^-^ parameters *vs* CD8^-^ function. For the DP thymocyte system, we fitted 11 biophysical parameters *vs* 1/EC_50_ plots of 5 (in the absence of CD8) or 7 (in the presence of CD8) data points by three functions: linear, exponential, and sigmoidal in each graph (Supplementary Fig. 5). For the SP naïve T cell system, we normalized the 4 functional readouts by scaling each by its maximum (*i*.*e*. the value for OVA) and by a negative control (i.e. the value for BSA), plotted them against biophysical parameters of OT1 TCR interacting with 3 peptides in the same graph, and fitted the resulting 12 points by three functions: linear, exponential, and sigmoidal (Supplementary Fig. 6)

The goodness-of-fit of the three functions was assessed by the *R*^2^ values. The correlation between biophysical parameters and functional readouts was further assessed by the non-parametric Spearman’s rank correlation coefficients *ρ* (which measures the strength and direction of association between two ranked variables) as well as the *p*-values for the significance of the Spearman’s correlation, all indicated in each panel (Supplementary Fig. 5 and 6). Using two heatmaps, one for the DP (Fig. 3A) and the other for the SP (Fig. 3B) system, we depict the 44 values for each of the four quantifiers – *ρ, p, x*_0_ and 1/*k* (*x*_0_ = midpoint and *k* = slope at *x*_0_ of the sigmoidal fits) – for the degrees of correlation of the 11 biophysical parameters, 4 quantifiers per biophysical parameter for the three cases of CD8^+^ *vs* CD8^+^, CD8^-^ *vs* CD8^+^, and CD8^-^ *vs* CD8^-^.

**Figure 3.**
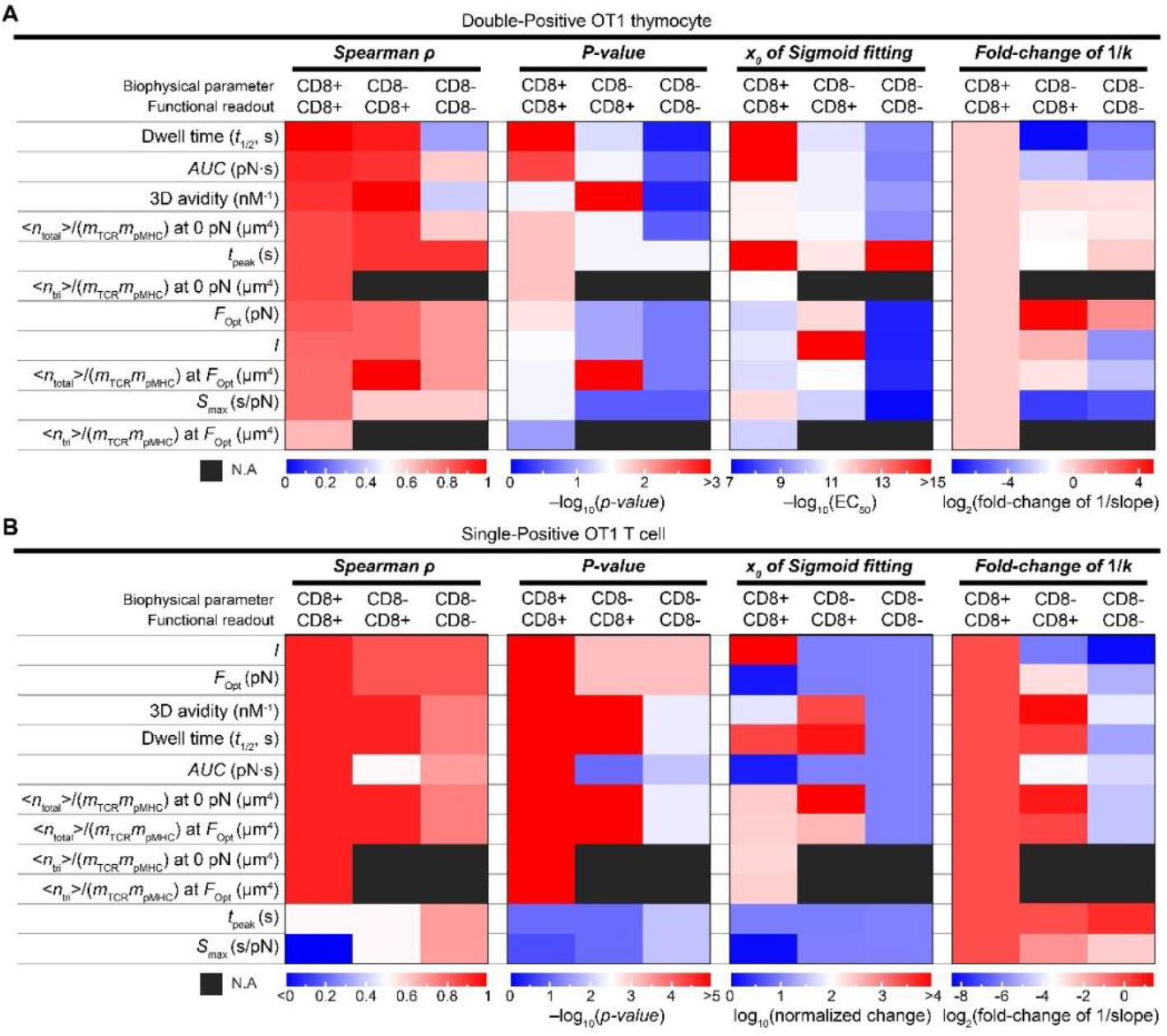
Correlating biophysical metrics with biological functions. **(A, B)** *1*^*st*^ *column*: List of 11 biophysical parameters measured by micropipette adhesion frequency, BFP force-clamp, and tetramer binding assays. *2*^*nd*^ *– 12*^*th*^ *columns*: four quantifiers (Spearman’s rank correlation coefficients *ρ*, the *P*-values for the significance of the Spearman’s correlation, *x*_0_ (midpoint of the sigmoidal fits), and fold-change of 1/*k* (*k* = slope at *x*_0_ of the sigmoidal fits)) for the three cases of CD8^+^ *vs* CD8^+^, CD8^-^ *vs* CD8^+^, and CD8^-^ *vs* CD8^-^, which represent three combinations of measurements made in the presence (+) and absence (-) of CD8: 1) CD8^+^ parameters *vs* CD8^+^ function, 2) CD8^-^ parameters *vs* CD8^+^ function, and 3) CD8^-^ parameters *vs* CD8^-^ function. All correlation analyses were performed using functional readout(s): CD69 upregulations for DP thymocytes ((A) and Supplementary Figure 5) and CD3, CD25, CD69, and Ca^2+^ *AUC* for the SP naïve T cells ((B) and Supplementary Figure 6).

Comparing the heatmap patterns in Fig. 3A and 3B reveals similar results from DP thymocytes and SP naïve T cells. By visualizing the progressive cooling of colors from red to blue when moving from the left column rightwards (Fig. 3), it is evident that the biophysical parameters correlate with biological responses the best when both were measured in the presence of CD8 (CD8+ *vs* CD8+) and the least when both were measured in the absence of CD8 (CD8- *vs* CD8-), with mid-level correlations between the biophysical parameters measured in the absence of CD8 and the biological responses measured in the presence of CD8. Specifically, the decreased power by the CD8^-^ biophysical parameters to predict the CD8^-^ (Supplementary Figs. 5A-5I and 6A-6I) relative to the CD8^+^ (Supplementary Figs. 5J-5R and 6J-6R) functions can be explained by the poor biological activities of the altered peptide ligands (APLs) when CD8 was absent. The increased power by the CD8^+^ (Supplementary Figs. 5S-5AC and 6S-6AC) relative to the CD8^-^ (Supplementary Figs. 5J-5R and 6J-6R) biophysical parameters to predict the CD8^+^ functions indicates that the biophysical metrics measured in the presence of CD8 contain additional information, which enables them to amplify the predictive power of the counterpart metrics measured in the absence of CD8.

By sorting the 11 biophysical parameters in each column, we obtain a ranking of their ability to predict the T cell response. However, three caveats are noted. First, for the DP thymocyte system, force-based and 2D data include 7 peptides: OVA, Q4, Q4R7, T4, Q4H7, Q7, and G4; whereas the 3D data includes only the first 5 of these. A smaller number of data points likely yields better Spearman coefficients for the 3D avidity *K*_v_ and Dwell time *t*_½_. Second, because of the low number of data points in these plots, many of the Spearman coefficients are of similarly high values. An extreme case is the SP naïve T cell system, where both biophysical parameters and biological functions were measured in the presence of CD8, with 9 of 11 metrics showing the same levels of correlation by 3 of the four quantifiers, as evidenced by the same red color in the CD8^+^ *vs* CD8^+^ columns of the *ρ, p, x*_0_ and 1/*k* groups (Fig. 3B). Third, while consistencies are observed among some biophysical parameters across some quantifiers, not all quantifiers predict the same ranking for all three combinations of measurements with and without CD8 (Fig. 3), thereby preventing us from pinpointing the best biophysical predictor for biological function.

### 3.5 CD8 enhances TCR specificity for ligands across threshold of thymocyte negative selection

In addition to inducing quantitatively different CD69 expressions on DP thymocytes (Fig. 3A), the 7 peptides behave qualitatively differently in the fetal thymic organ culture (FTOC) assay to be distinctively classified as negative selection (OVA, Q4, Q4R7), threshold (T4), and positive selection (Q4H7, Q7, G4) ligands (29, 70). The importance of CD8 in this thymocyte negative selection process was suggested by the peptide scanning model (29). It was further supported by our finding of two concurrent changes in the biophysical metrics and in the cell fates when compared thymocytes expressing WT CD8 or a chimeric molecule CD8.4 (the CD8 ectodomain fused with the CD4 cytoplasmic domain to increase Lck association (71)). The cell fate change was observed in the FTOC experiment in which the negative selection threshold was shifted rightward from T4 to Q4H7 (29). The change in the biophysical metrics was observed in the BFP experiment in which the demarcation between coreceptor capable *vs* incapable of cooperating with TCR in pMHC binding to form a tripartite dynamic catch was shifted rightwards from Q4R7 to Q4H7 (22).

Therefore, we next added the cell fate outcomes in thymocyte negative selection as a functional readout to correlate with the biophysical parameters. To build upon our previous analysis (22), we plotted the percentage of CD8^+^ SP thymocytes that survived the FTOC assay *vs* the four 2D affinity-based parameters (Fig. 4A-4D) and the five bond profile metrics (Fig. 4E-4I) for both cases of without (gray open circle) and with (black close circle) CD8. The negative selection threshold can be identified from the sharp drop of the CD8^+^ SP thymocyte survival levels from >50% to <10% for different peptides (indicated) by dashed vertical lines. The ranges of parameter change (i.e., the difference between the same parameter evaluated using Q4R7 and Q4H7) are marked by purple (without CD8) and red (with CD8) vertical stripes.

**Figure 4.**
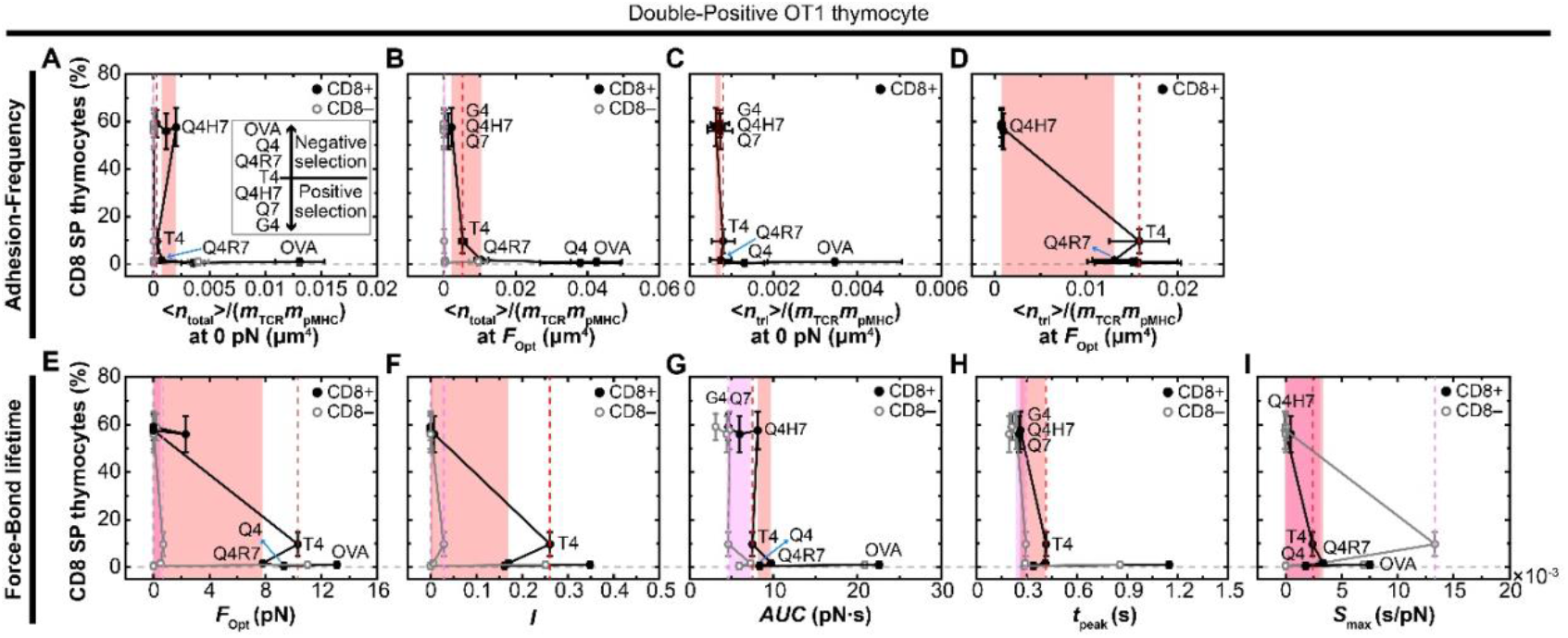
CD8 impacts on the TCR specificity for two ligands evaluated by 2D measures. **(A-I)** The percentage of CD8^+^ SP thymocytes from FTOC assay are plotted *vs* biophysical metrics of OT1 TCR interacting with a panel of 7 peptides (OVA, Q4, Q4R7, T4, Q4H7, Q7, G4) presented by H2-K^b^ (CD8^+^, closed black circles connected by black line segments) or 5 peptides (OVA, Q4R7, T4, Q4H7, G4) presented by H2-K^b^α3A2 (CD8^-^, open gray circles connected by gray line segments) to permit or prevent CD8 from binding to MHC measured from the adhesion frequency assay and force-clamp assay by the BFP: normalized average number of bonds at zero force 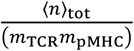 (A) and at optimal force 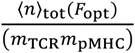 (B), normalized synergy at zero force 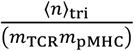 (C) and at optimal force 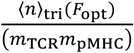 (D), optimal force *F*_opt_ (E), catch bond intensity *I* (F), area under the bond profile curve A*UC* (G), peak bond lifetime *t*_peak_ (H), and the maximum slope along the curve *S*_max_ (I). Vertical dashed lines identify the parameters of the threshold peptide (T4) and the vertical stripes mark the parameter ranges across the threshold from the strongest positive selection peptide (Q4H7) to the weakest negative selection peptide (Q4R7) using different colors to indicate measurements made when CD8 was prevented (purple) or permitted (red) to bind MHC. The FTOC assay data are directly from (70). The 2D binding parameters are either directly taken from (22) ( 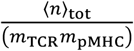 and 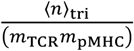) or calculated ( 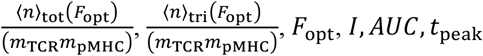, and *S*_max_) by fitting the data from (22) using the two-pathway model (cf. Supplementary Figure 3). Error bars are directly from experimental data from published data from (22, 70).

It is evident from the narrow purple stripes that except for *AUC*, all other parameters do very poor jobs to distinguish between negative *vs* positive selection ligands in the absence of CD8 (Fig. 4). CD8 has a small impact on the ability of the force-free 2D-based parameters 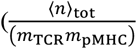 and 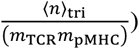 and 3D-based parameters (dwell time *t*_1/2_ and avidity *K*_v_) to differentiate two peptides, Q4H7 (the strongest positive selection ligand) and Q4R7 (the weakest negative selection ligand), that reside on the two sides of the negative selection threshold (Fig. 4A, C and Supplementary Fig. 7). Based on the changes in the widths of the purple stripes from that of the red stripes, CD8 seems to have amplified this ability moderately for the force-free 2D-based parameters at optimal force (Fig. 4B and D). It was also observed that the amplified ability by CD8 becomes obvious for both the optimal force (Fig. 4E) and the catch bond intensity (Fig. 4F), negatively for the *AUC* (Fig. 4G), mildly for both peak bond lifetime (Fig. 4H) and the maximum slope (Fig. 4I). However, this may not be precise because the different parameters have different units, making direct comparison of their absolute changes among different parameters difficult.

To overcome this difficulty, we normalized the ranges of parameter change using the maximum value of that parameter measured using OVA:H2-K^b^. We plotted in Fig. 5A the relative changes for the nine 2D parameters and two 3D parameters depicted in Fig. 4 and Supplementary Fig. 7 in the absence and presence of CD8. Except for *AUC* and *S*_max_, no other parameters measured in the absence of CD8 show appreciable relative changes across the negative selection threshold (Fig. 5A), consistent with the impression from comparing the non-normalized cross-threshold changes (Fig. 4). In the presence of CD8, 2D parameters (normalized average number of total bonds and normalized synergy) measured at zero force show negligible increase in values across the threshold (Fig. 5A). Interestingly, when measured under force, all 2D parameters show appreciable relative changes, except for *AUC*. Of these, the optimal force, the catch bond intensity, the maximum slope, the normalized average number of bonds, and the normalized synergy show similar relative increases. These high levels (>40 %) of relative changes suggest that broadening the catch bond force regime is more important than prolonging the overall bond lifetime for the discrimination of negative and positive ligands.

**Figure 5.**
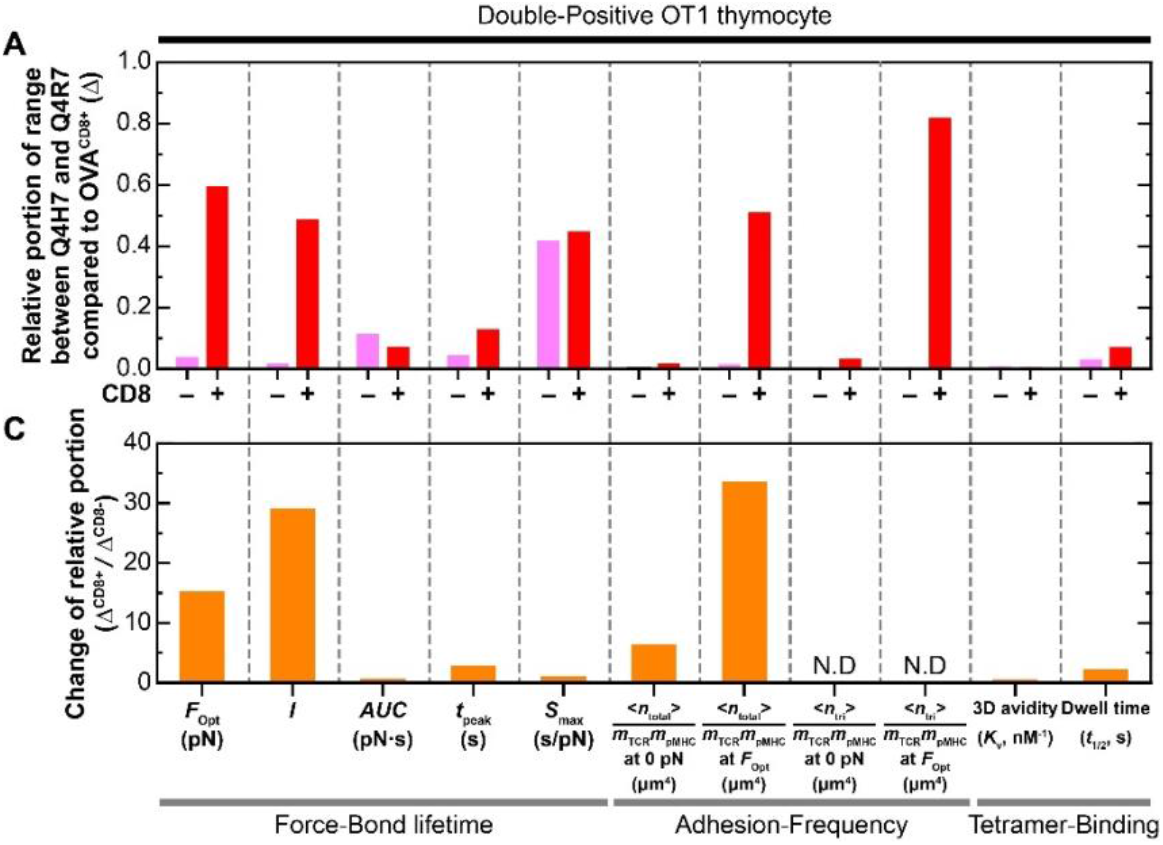
TCR specificity as measured by biophysical parameters. **(A)**, Change of parameters (the same as those in Figure 4 and Supplementary Figure 7) across the negative selection threshold between values measured for OT1 TCR binding to Q4H7 and Q4R7 presented by H2-K^b^α3H2 (Δ^CD8-^ = Q4R7^CD8-^ – Q4H7^CD8-^, purple bar) and H2-K^b^ (Δ^CD8+^ = Q4R7^CD8+^ – Q4H7^CD8+^, red bar) normalized by that to OVA presented by H2-K^b^ (OVA^CD8+^, the maximum value). **(B)**, Ratios of changes (Δ^CD8+^/ Δ^CD8-^) for the indicated paired parameters.

We also plotted the ratios of these relative changes measured in the presence over the absence of CD8 in Fig. 5B. This ratio quantifies how much the range of difference of the OT1 TCR interaction parameters with Q4R7 and Q4H7 is widened by permitting CD8 to bind MHC relative to preventing CD8 from binding. It can be seen from Fig. 5B that, of the nine 2D parameters analyzed, three force-based parameters (optimal force, the catch bond intensity, and the normalized average number of bonds evaluated by the optimal force) show high levels of amplifications. In other words, CD8 and force enhance the specificity of TCR for the two peptides across the threshold of thymocyte negative selection.

By comparison, 3D parameters (*e*.*g*., affinity/avidity and dwell time) which do not contain any information of force-response, show extremely narrow stripes across the negative selection threshold regardless of whether CD8 is present or absent (Supplementary Fig. 7). For 3D avidity, the normalized ranges of cross-threshold change measured in the presence and absence of CD8, and their ratio are all negligibly small (Fig. 5A). For 3D dwell time, the normalized ranges of change are larger, especially when the measurements were made in the presence of CD8, resulting in a small but non-negligible amplification factor of 2 by the CD8 (Fig. 5B). Consequently, 3D *K*_v_ and *t*_1/2_ show negligible values in Fig. 5.

### 3.6 The effect of CD4 in class II restricted TCR system

Compared to CD8, CD4 binding to pMHC-II is crucial for T cell functions such as regulation and suppression of immune reactions as well as communicating and activating various immune and non-immune cells (72-76). However, studying the role of CD4 binding has been difficult due to ultra-low affinity of CD4 for pMHC-II (20). Notwithstanding the very small amount of available data for TCR–pMHC-II–CD4 trimolecular interactions, we nevertheless tested whether the biophysical metrics of bond profiles can also be used to discern the changes between TCR-mediated T cell activation in the absence and presence of CD4. We re-analyzed 5 pairs of force-dependent bond lifetime curves measured by the Zhu lab using BFP. One pair is for the E8 TCR interacting with TPI:HLA-D1 in the absence and presence of CD4 (20) (Supplementary Fig. 2A). Four pairs are for the murine 3.L2 TCR expressed on either CD4^+^CD8^-^ or CD4^-^CD8^+^ SP T cells interacting with 4 peptides (Hb, T72, I72, and A72) bound to I-E^k^ (43, 53) (Supplementary Fig. 3G). Similar to the MHC class I restricted TCR case (Fig. 2A), in the case of MHC class II restricted TCR, permitting *vs* preventing CD4 from cooperating also increased some of the five metrices of bond profiles for some interactions, as seen by the cool to warm color changes from the CD4- to CD4+ columns in Fig. 6A (see also Supplementary Fig. 2B). Unlike the MHC class I restricted TCR case (Fig. 3), in the case of MHC class II restricted TCR, data are available for correlations of only two pairs: 1) CD4^+^ parameters *vs* CD4^+^ function, 2) CD4^-^ parameters *vs* CD4^+^ function. Nevertheless, our 2D biophysical parameters showed general correlations with ligand potency which are enhanced by measuring the biophysical parameters in the presence of CD4, although the enhancements are not always evident (Fig. 6B and Supplementary Fig. 8). Additionally, similar to the case of MHC-I, we found that the sigmoid fitting in biophysical parameter *vs* function domain yields the best fit, underscoring the non-linearity of the relationship between biophysical parameters and T cell function.

**Figure 6.**
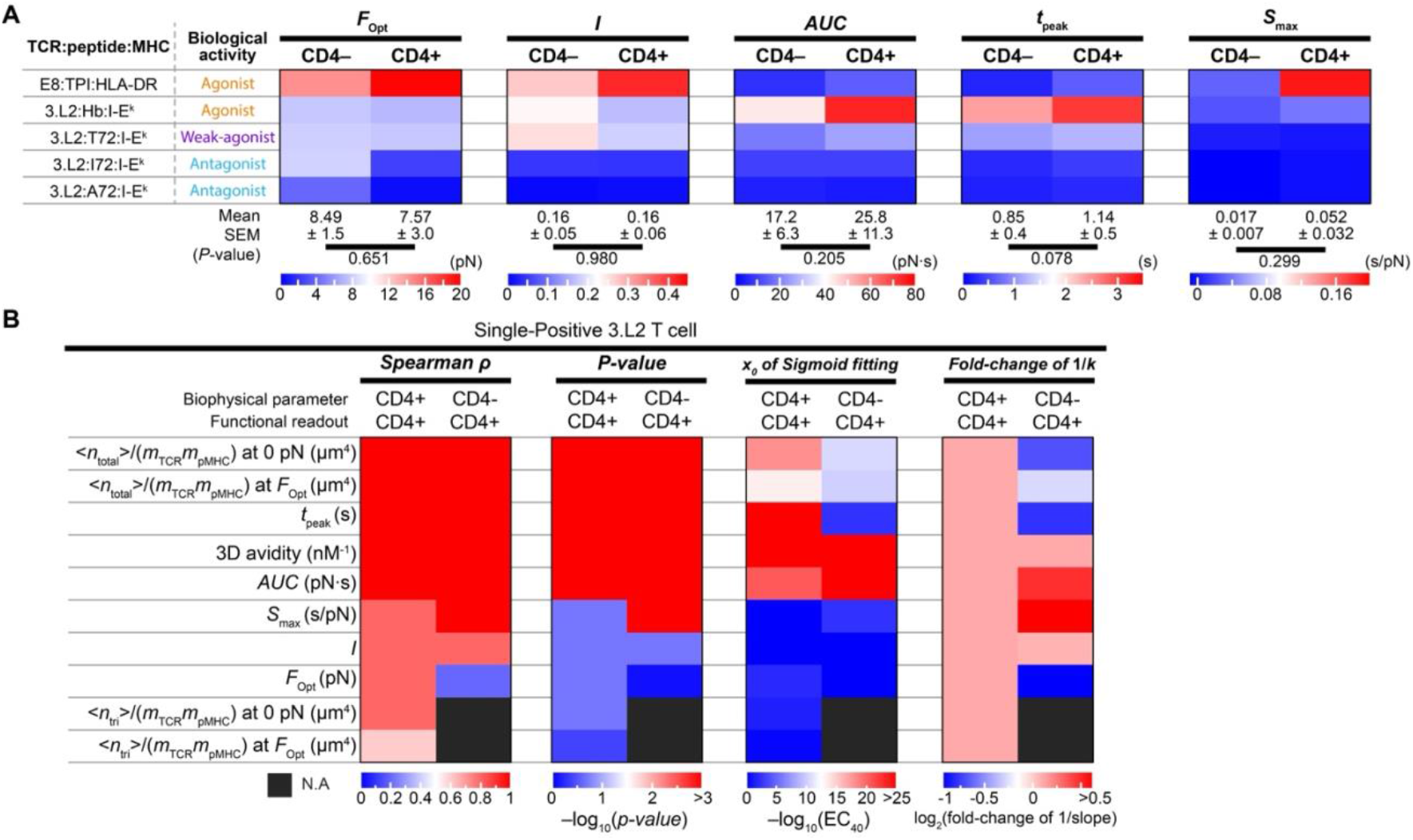
Evaluating metrics of TCR bond profiles without and with CD4 cooperation and their correlation with biological activity. **(A)** *1*^*st*^ *column*: List of 5 pairs of interactions between TCR and pMHC-II. *2*^*nd*^ *column*: Biological activities of these interactions based on literature, which are grouped into three color-coded categories (agonist = orange, weak agonist = purple, antagonist = cyan). *3*^*rd*^ *– 12*^*th*^ *columns*: Five pairs of metrics (***F***_**opt**_, ***I, AUC, t***_**peak**_, **and *S***_**max**_, as indicated on the top of the 1^st^ row) of force-lifetime curves of 2 TCRs forming catch and slip bonds with their respective panels of pMHCs without (CD4-) and with (CD4+) coreceptor cooperation. Their values are shown as heatmaps with specific color codes for each metric indicated at the bottom. For each interaction pair, the change in color indicates the change in the metric values between the case in which CD4 was prevented from binding pMHC and the case in which CD4 was permitted to bind pMHC. The mean and SEM values across all 5 pairs of interactions are also shown in the bottom, as well as the *P*-values that indicate the significance of the increases in the values from the CD4-column to the CD4+ column. **(B)** *1*^*st*^ *column*: List of 10 biophysical parameters measured by micropipette adhesion frequency, BFP force-clamp, and surface plasmon resonance (SPR). *2*^*nd*^ *– 9*^*th*^ *columns*: four quantifiers (Spearman’s rank correlation coefficients *ρ*, the *P*-values for the significance of the Spearman’s correlation, ***x***_**0**_ (midpoint of the sigmoidal fits) and fold-change of **1**/***k*** (***k*** = slope at ***x***_**0**_ of the sigmoidal fits)) for the two cases of CD4^+^ *vs* CD4^+^ and CD4^-^ *vs* CD4^+^. All correlation analysis were performed using the logarithm of the reciprocal peptide concentration required to stimulate 40% maximal IL2 (1/EC40) as functional readout(s) ((A) and Supplementary Figure 8).

## 4 Discussion

The goals of this work are two-fold: 1) to examine the correlation of (or the lack thereof) TCR catch and slip bonds with T cell function and 2) to examine the concomitant prolongation of (or the lack thereof) force-dependent TCR bond lifetime and the corresponding enhancement of T cell function, both from the situation when the coreceptors CD8 and CD4 are permitted to bind pMHC-I/II concurrently to the situation when they are prevented from binding. To evaluate the correlation of TCR catch bonds with their agonist ligands and slip bonds with their antagonist ligands, originally suggested in previous publications (22, 42, 43, 45-48), the present study has analyzed a much larger dataset (26 pairs of bond lifetime *vs* force curves), used multiple quantitative metrics for bond profiles instead of qualitative description, and included data measured not only from the situation when CD8/CD4 are permitted to bind pMHC-I/II but also from the situation when CD8/CD4 are prevented from binding. We found that such correlation works well using data obtained from the same system but less well across different systems. For example, the 2C TCR forms catch bond, slip bond, and slip bond with three peptides (R4, dEV8, and EVSV) presented by H2-K^b^α3A2 or H2-K^b^ (Supplementary Fig. 3A) but functional measurements suggest that R4 behaved as super agonist, dEV8 as weak agonist, and EVSV as antagonist (46). By comparison, biophysical measurements show that the P14 TCR forms progressively weaker and weaker catch bonds with three gp33 peptides (41M, 41C, and 41CGI) (Supplementary Fig. 3D) but functional measurements suggest that 41M behaved as super agonist, 41C as agonist, and 41CGI as weak agonist (48). Yet, the force-lifetime curves of the P14 bond with 41CGI and the 2C bond with dEV8 are quite different regardless of whether CD8 was prevented or permitted to bind. Still, the OT1 TCR forms a catch bond with the strong agonist peptide OVA that looks very similar to the P14 catch bond with 41CGI when both measured in the presence CD8. In addition, the N15 TCR catch bonds with VSV8 in the absence or presence of CD8 are much more pronounced with lifetimes much longer than any other interaction pairs analyzed in this study (Supplementary Fig. 3F). Despite these differences, our UMAP analysis, which takes into considerations five metrics of force-lifetime curves both with and without CD8 contributions, groups interactions of N15 with VSV8, OT1 with OVA, 2C with R4, and P14 with all three peptides (41M, 41C, and 41CGI) into cluster 1, all members of which are agonists except for the 41CGI case (Fig. 2B).

Besides the differences in TCR systems, other causes of these inconsistencies may include employing different experimental techniques (BFP *vs* optical tweezers) and using TCRαβ ectodomain proteins *vs* cell surface TCR-CD3 complex, as mentioned earlier. To examine whether using living T cells for TCR–pMHC bond lifetime measurements, which is known to induce intracellular calcium fluxes (42, 77), would impact the measured values, we evaluated the memory index, which measures the increase in the likelihood of TCR to bind pMHC in a given contact in a series of repeated contacts between a T cell and an APC, given that the immediate prior contact resulted in binding relative to no binding (51, 78). Building upon our previous observation of the memory effect from the adhesion frequency between T cell and APC in the absence of CD8 (51, 78), in this study we showed such memory in the presence of CD8 (Supplementary Fig. 9A-D). Our previous observation also suggests that such memory effect is limited to on-rate, as lifetimes measured from bonds with or without an immediate prior binding or lifetime event are indistinguishable and that repeatedly exerting durable forces on TCR–pMHC bonds during the process of measuring their force-dependent lifetimes would not change the bond lifetime over time, despite that the T cell was activated by such stimulations (78). Adding to these previous findings for the case when CD8 was prevented from binding, the present work observed the same results for the case when CD8 was permitted to bind (Supplementary Fig. 9E-H). Whether, and if so, how such memory index as a biophysical metric may serve as a predictor for T cell activation and function should be an interesting topic for future studies.

Two types of biophysical metrics have been used as predictors of T cell function. One is binding affinity. The rationale is that the higher the affinity, the greater the TCR bond numbers to trigger T cell activation, giving the same densities of TCR and pMHC. The other is bond lifetime. The rationale stems from the original kinetic proofreading model (32), for the longer the TCR engagement with ligand, the further the downstream signaling processes (protein docking and conformational changes, enzymatic modifications, formation of molecular assemblies, *etc*.) may proceed or the greater chance for these processes to reach a point of no return upon TCR disengagement (49, 79). A modification of the kinetic proofreading model has incorporated the idea of rebinding (80), providing the rationale for using biophysical metrics that combine both affinity and bond lifetime. Since TCR may experience forces that alter their bond lifetime with ligand, a question arises as to which lifetime duration at what force level should be used to relate to TCR signaling. For TCR interactions that form slip bonds, *i*.*e*., those with antagonist or weak agonist ligands, their bond lifetimes reach the longest values at zero force; and the choice of *t*_0_ as the T cell functon predictor is consistent with the use of 3D dwell time (*t*_1/2_) in the original form of the kinetic proofreading model that does not consider the effect of force (32). In contrast, for TCR interactions that form catch bonds, *i*.*e*., those with agonist ligands, their bond lifetimes reach the longest value *t*_peak_ at the optimal force *F*_opt_ (> 0, *F*_opt_ = 0 for slip bonds), and are longer than the force-free value *t*_0_ over a range of forces 0 < *F*< *F*_range_ (Fig. 1B). It is interesting (but not yet understood) that data measured using BFP and optical tweezers thus far all report *F*_opt_ and *F*_range_ values to be around 10-20 pN and 20-40 pN, respectively, for agonists, regardless of whether the peptide is presented by MHC-I or II. Perhaps it is not coincidental that newly developed DNA origami tension sensors showed that TCRs transmit 10–20 pN forces to antigens (81), which confirms the previous results of the same lab obtained using an earlier version of the DNA tension probe (82). However, this is inconsistent with the results of much smaller forces (∼2 pN) obtained by another group using a spider sick peptide based tension probe (83). Also inconsistent with the catch bond profile of cell surface 1G4 TCR measured using BFP in the 0-22 pN force range (46) is the slip bond profiles of soluble 1G4 TCRαβ ectodomain proteins measured using a flow chamber in forces >10 pN (84). These authors also suggest 3D affinity and dwell time of TCR–pMHC bond to be predictors of T cell activation and parameters for antigen discrimination (85). Consequentally, they suggest that force may impair antigen discrimination by reducing differences in dwell time (84).

In the present study, we examined correlations of (and the lack thereof) of T cell responses with five biophysical metrics of the bond profiles (*t*_peak_, *F*_opt_, *I, AUC*, and *S*_max_) across eight MHC-I restricted TCR systems and two MHC-II restricted TCR systems. We mainly compared the predictive powers of these metrics with two 2D metrics measured at zero force, 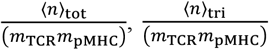, and with the same two parameters calculated at optimal force. Using data available from the thymocyte negative selection studies, we also compared these with two 3D binding parameters, *K*_v_ and dwell time (*t*_1/2_). These correlative analyses suggest that force-based metrics better predict T cell function and discriminate antigen, supporting the view that mechanical forces usually exert on TCR and such forces are important, as they can amplify TCR antigen discrimination as well as T cell signaling and function, regardless of whether force is required to trigger TCR in every and all circumstances. Recent steered molecular dynamics (SMD) simulations found that mechanical stability metrics of TCR–pMHC bonds (peak rupture force and mechanical work of unbinding) strongly correlate with signaling events and cell function (86), providing a computational counterpart to the experimentally determined force-lifetime metrics presented here, which corroborates with the idea that force-based parameters are more appropriate predictors of T cell function. Future studies will be required to reconcile the apparent discrepancies between these results and those of Pettmann *et al*. (84, 85).

Another disagreement between us and Pettmann *et al*. is how to measure the TCR antigen discrimination power. Pettmann *et al*. quantify the discrimination power as the ability of TCR to amplify small changes in pMHC affinity into larger changes in ligand potency, seemingly independent of their definition of antigen sensitivity, which quantifies the lowest pMHC affinity capable of triggering TCR signaling (85). In comparison, we used the thymocyte selection data to perform a detailed analysis to reveal the two competing requirements for antigen discrimination by the TCR – sensitivity and specificity – which must be taken into account simultaneously, hence involving a tradeoff. On the one hand, evaluating by functional outcome, discrimination requires dually that thymocytes undergo apoptosis when they interact with Q4R7, *i*.*e*., to generate a true positive response when they should (sensitivity), and that thymocytes survive when they interact with Q4H7, *i*.*e*., not to generate a false positive when they should not (specificity). The larger the differential functional outcomes (Fig. 4, *y*-axis values of percent survival of CD8^+^ SP cells from the FTOC assay), the higher the specificity, and the greater the discriminatory power. On the other hand, evaluating by TCR recognition, discrimination requires dually that the thymocytes exhibit above threshold binding parameters when they interact with Q4R7, *i*.*e*., to generate a true positive response when they should (sensitivity), and that the thymocytes exhibit below threshold binding parameters when they interact with Q4H7, *i*.*e*., not to generate a false positive when it should not (specificity). The larger the differential binding parameters (Fig. 4, the widths of the colored strips), the higher the specificity, and the greater the discriminatory power.

Given that the OT1 thymocytes can discriminate between Q4R7 and Q4H7 to result in distinctive functional outcomes (29, 70), we asked what biophysical metric(s) of TCR interaction with ligand can best capture the mechanism. We hypothesized that this biophysical metric(s) may be identified by its greatest differential values between Q4R7 and Q4H7 to allow for the highest specificity, as the larger the differential values, the smaller the chance for stochasticity of individual measurements to generate a false positive. We therefore compared the ranges of changes of these metrics across the thymocyte negative selection threshold to rank their importance to the TCR specificity. To evaluate the effects of force and CD8 on the TCR specificity for peptides across the threshold of thymocyte negative selection, we further examined whether, and if so, how much such ranges would be widened in the presence compared to the absence of force and when CD8 was permitted to bind compared to when CD8 was prevented from binding. We found that, for most cases, the parameters evaluated under force increase their values when CD8 is permitted, rather than prevented, from binding pMHC-I, consistent with the contention that CD8 binding prolongs TCR engagement, thereby enhancing T cell signaling. In general, we found that the metrics evaluated under force are more predictive than the force-free parameters, and the 2D parameters are more predictive than the 3D parameters, consistent with our previous findings (22) and views (79, 87, 88). These findings have provided further support to the proposal that catch bond measurements capture some aspects of the TCR mechanotransductive machinery that likely underpin its remarkable features of sensitivity, specificity, and ability to discriminate self *vs* nonself peptide, features that make the TCR to stand out among other cell surface receptors, but still stand defiant to reveal to us a clear mechanistic explanation. Our analysis of OT1 thymocyte negative selection complements a recent work (21) showing that the effect of CD8 cooperation on catch bonds is most informative within the medium force range (9-12 pN) and that lifetimes in this regime best correlate with T cell specificity. Notwithstanding that we used curve-level metrics that account for bond lifetimes at all forces, their conclusion that optimal catch bonds tuned by CD8 underlies antigen discrimination is fully consistent with our findings that force-based CD8-dependent metrics are the best predictors of biological activity across many TCR–pMHC pairs and both class I and II systems.

To decipher putative relationships between binding parameters and cellular responses, conventional approach primarily plots individual biophysical features of TCR bonds against T cell functional readouts to evaluate the quality of the correlation (36, 47, 50, 89). Although this approach remains useful and was used in this work for the sake of completeness (Supplementary Figs. 5, 6, and 8), it becomes increasingly constrained when the dataset spans numerous parameters, and when distinct biophysical parameters do not necessarily exhibit a unified trend. To overcome these difficulties, we took a data science approach to compare the quality of the fitting using a correlation-matrix format and to reduce the high dimensional data using UMAP analysis (Figs. 3 and 6). Not only does this provide a compact and unbiased depiction of how each biophysical property relates to diverse cellular outcomes, but it also reveals overall trends by consolidating large and diverse datasets. Thus, the conclusions we draw reflect the composite structure of the dataset rather than the behavior of any single parameter pair, allowing a more rigorous evaluation of how TCR mechanotransduction features integrate to regulate immune responses.

The evaluation of TCR catch bond properties and their correspondence to T cell activation and function is of interest in the context of adoptive cell therapy (ACT). In ACT, modified T cells expressing tumor antigen-specific TCRs are transferred to patients for immunotherapy against cancer (90, 91). A key issue is the selection of the most efficacious TCR(s) specific to neoantigen(s) and tumor associated antigen(s) of particular patients. As discussed earlier, three sets of parameters may be used: metrics evaluated under force, force-free 2D parameters, and 3D parameters. Whereas 3D affinity has been commonly used for evaluating TCR effectiveness, studies have shown an imperfect correlation between affinity and potency, as very high TCR affinities can lead to impaired T cell function with diminished antigen sensitivity *in vitro* and *in vivo* (92-99). Zhao et al. screened a library of mutated TCRs with altered complementary determinant regions (CDRs) and selected ones that had high signaling but moderate affinities (47). The authors found that those TCRs also form catch bonds with the pMHC in the presence of CD8, and the peak bond lifetime *t*_peak_ correlated well with CD69 upregulation (47).

In summary, this work showed that metrics derived from the force-lifetime curves of TCR bonds are useful predictors of T cell function in the case when the CD8/CD4 are permitted to bind pMHC-I/II, extending our previous results found when the coreceptors are prevented from binding. Our results provide better understanding of TCR catch bonds, shed new lights on the role of coreceptors, and support the TCR mechanotransduction hypothesis (49, 79).

## 5 Data Availability Statement

Other than data showing in Supplemental Figs. 2C (except for the blue curve in the left panel) and 4, which were generated in this work, original data supporting the findings of this study are from the cited publications and Jinsung Hong’s Ph.D. thesis available at Georgia Institute of Technology Library. The bond lifetime datasets re-analyzed for model fitting are summarized and made available at Zenodo (https://doi.org/10.5281/zenodo.11368090).

## 6 Ethics Statement

All experiments in this study were conducted following the protocols approved by the Institutional Review Board (IRB protocol No. H22021) and Institutional Care and Use Committee (IACUC protocol No. A100168) of the Georgia Institute of Technology.

## 7 Author Contributions

Conceptualization: S.T., H.-K.C., C.Z.; Methodology, S.T., H.-K.C., C.Z.; Investigation: S.T., H.-K.C., C.Z., A.H.K.A., M.L., V.E.W., P.C., L.D.; Writing: Original Draft-H.-K.C., C.Z.; Review & Editing: S.T., H.-K.C., C.Z., A.H.K.A., M.L., V.E.W., P.C., L.D.; Funding Acquisition: H.-K.C., C.Z.; Project Administration: H.-K.C., C.Z.

## 9 Acknowledgments

We thank Saurabh Sinha for discussion on methods of correlative analysis and NIH Tetramer Core Facility at Emory University for providing pMHC monomers used in this work. The original manuscript was submitted to *Scientific Reports* as a contribution to the Special Collection on “Immune Cell Receptor Signaling”. The present submission is a substantially revised version based on the reviewers’ comments, which we acknowledge.

## 11 Generative AI Statement

The authors declare that no Generative AI was used in the creation of this manuscript.

---

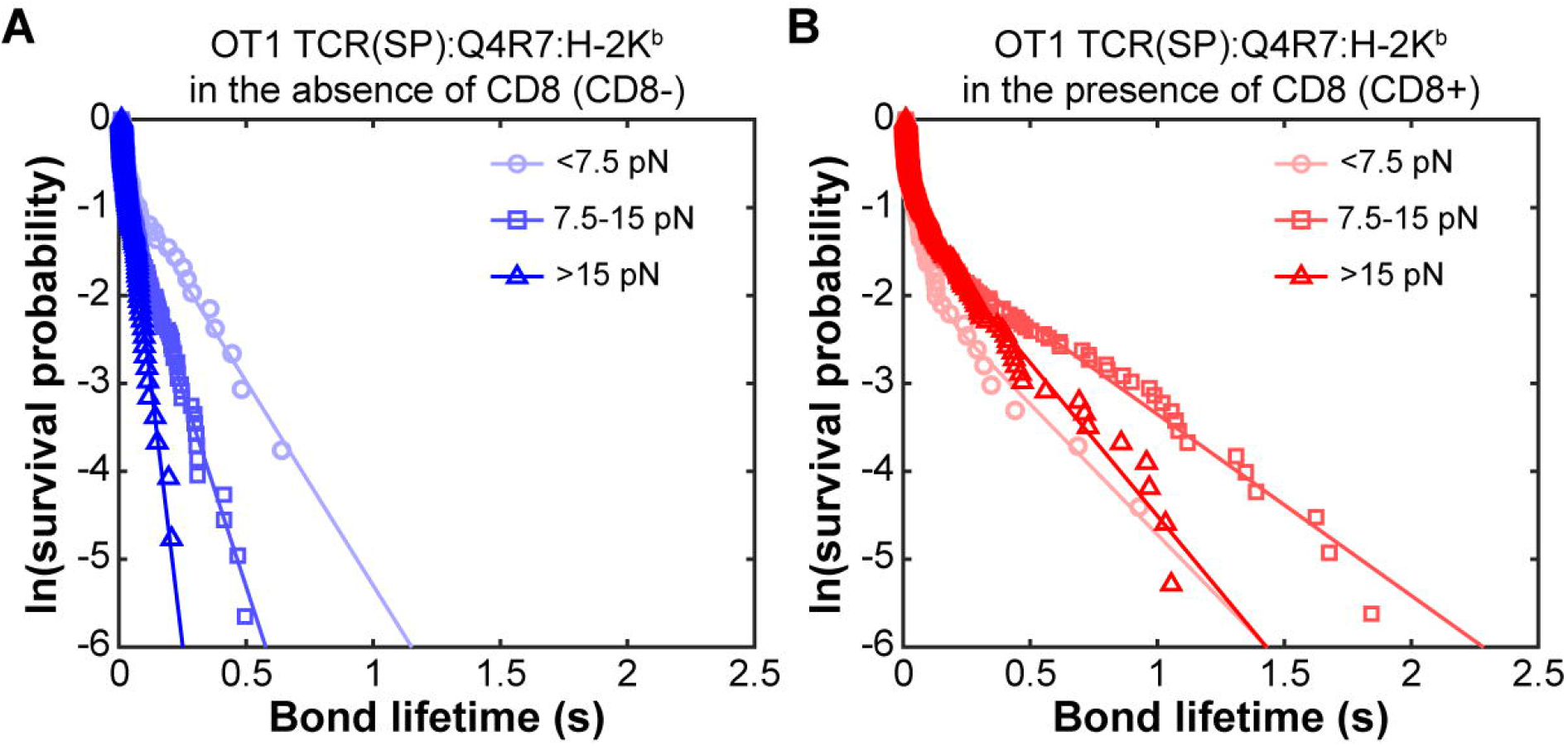

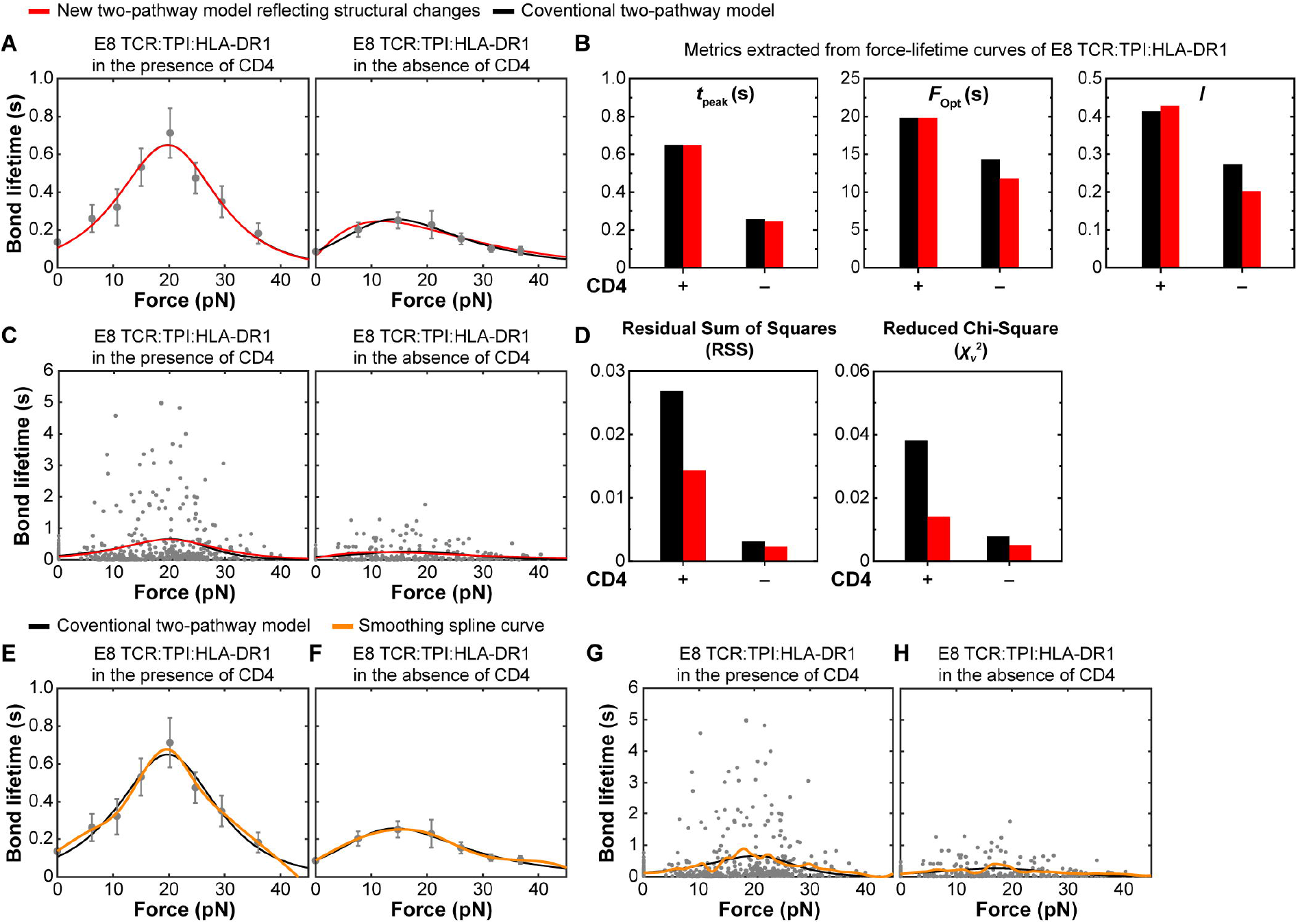

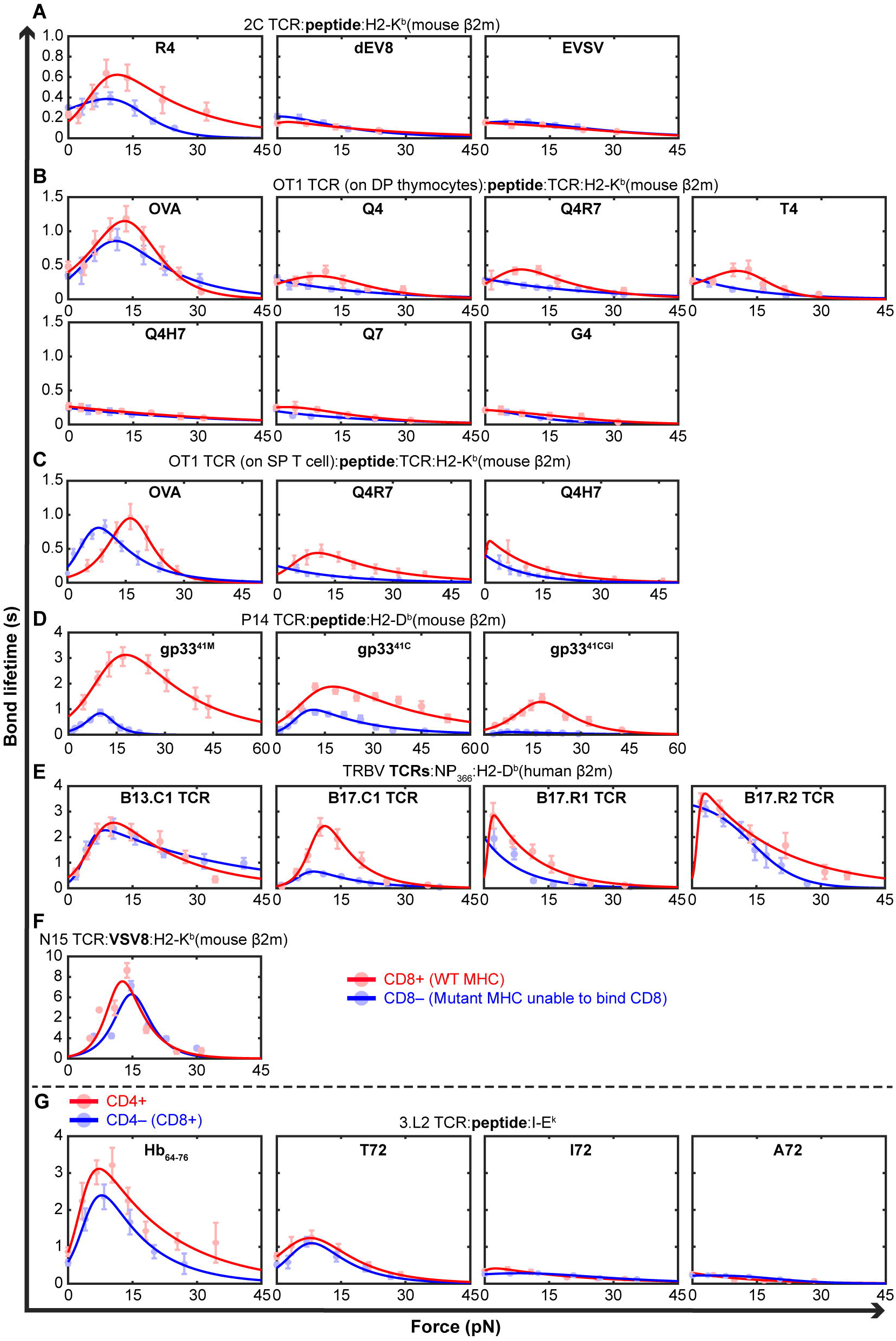

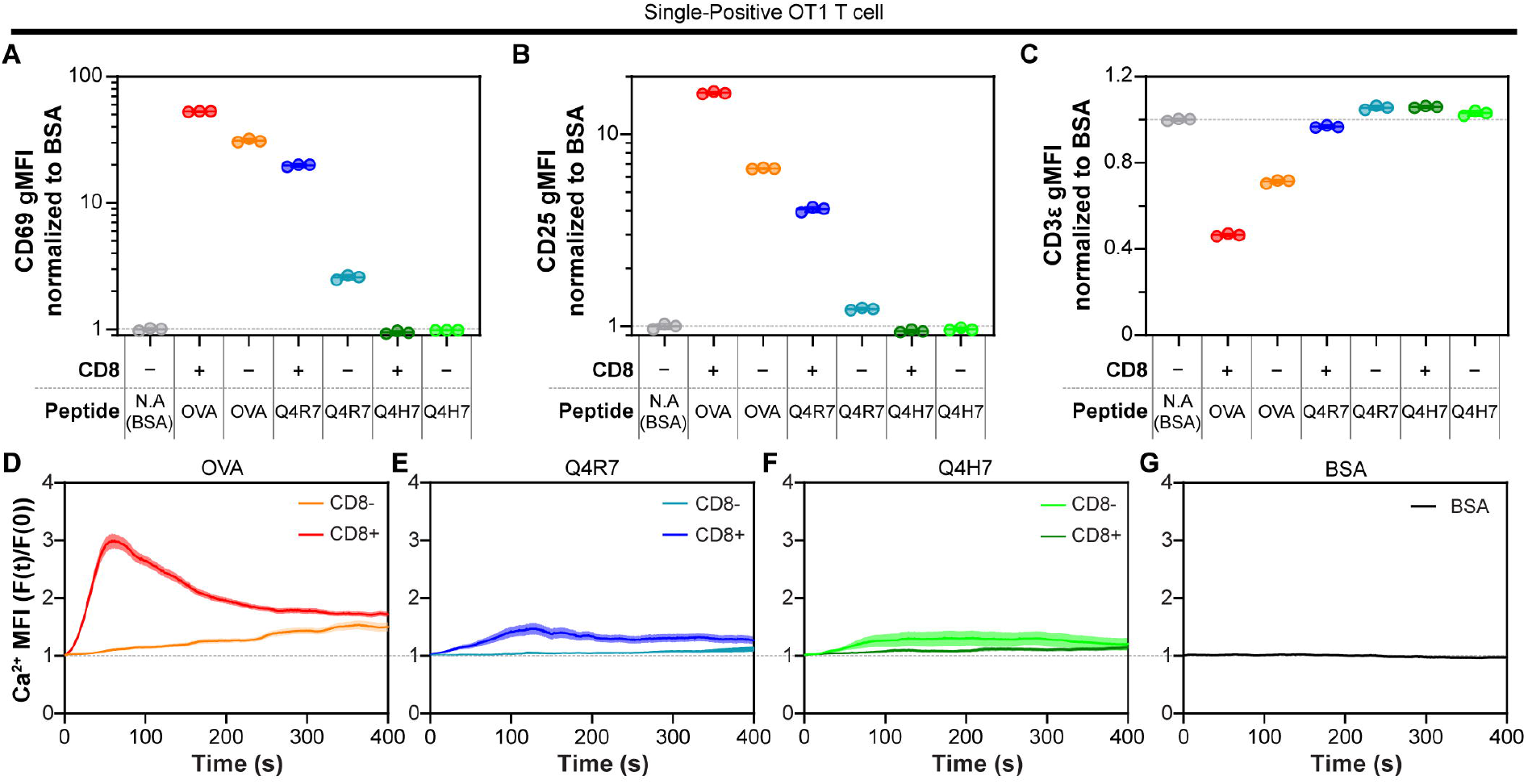

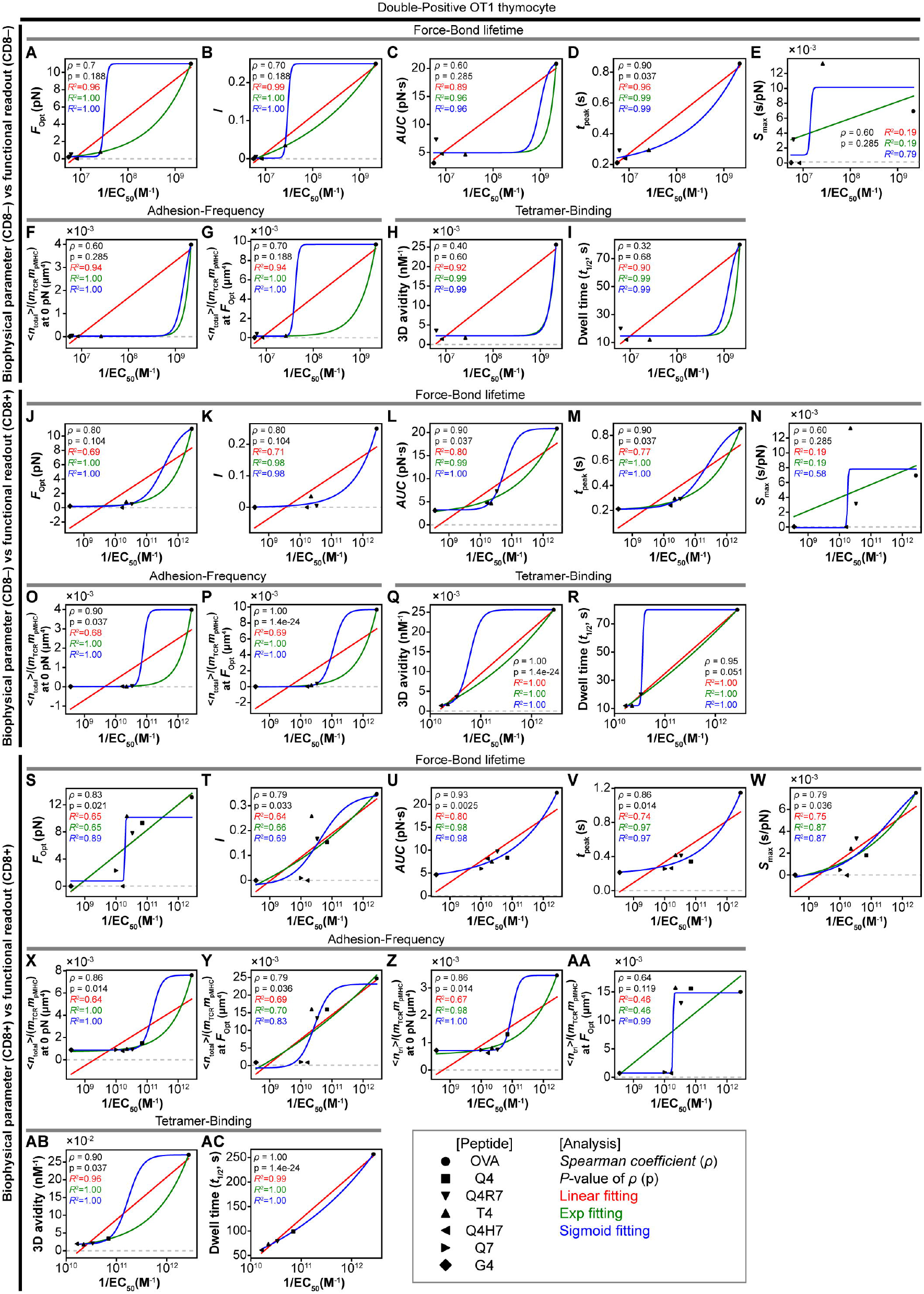

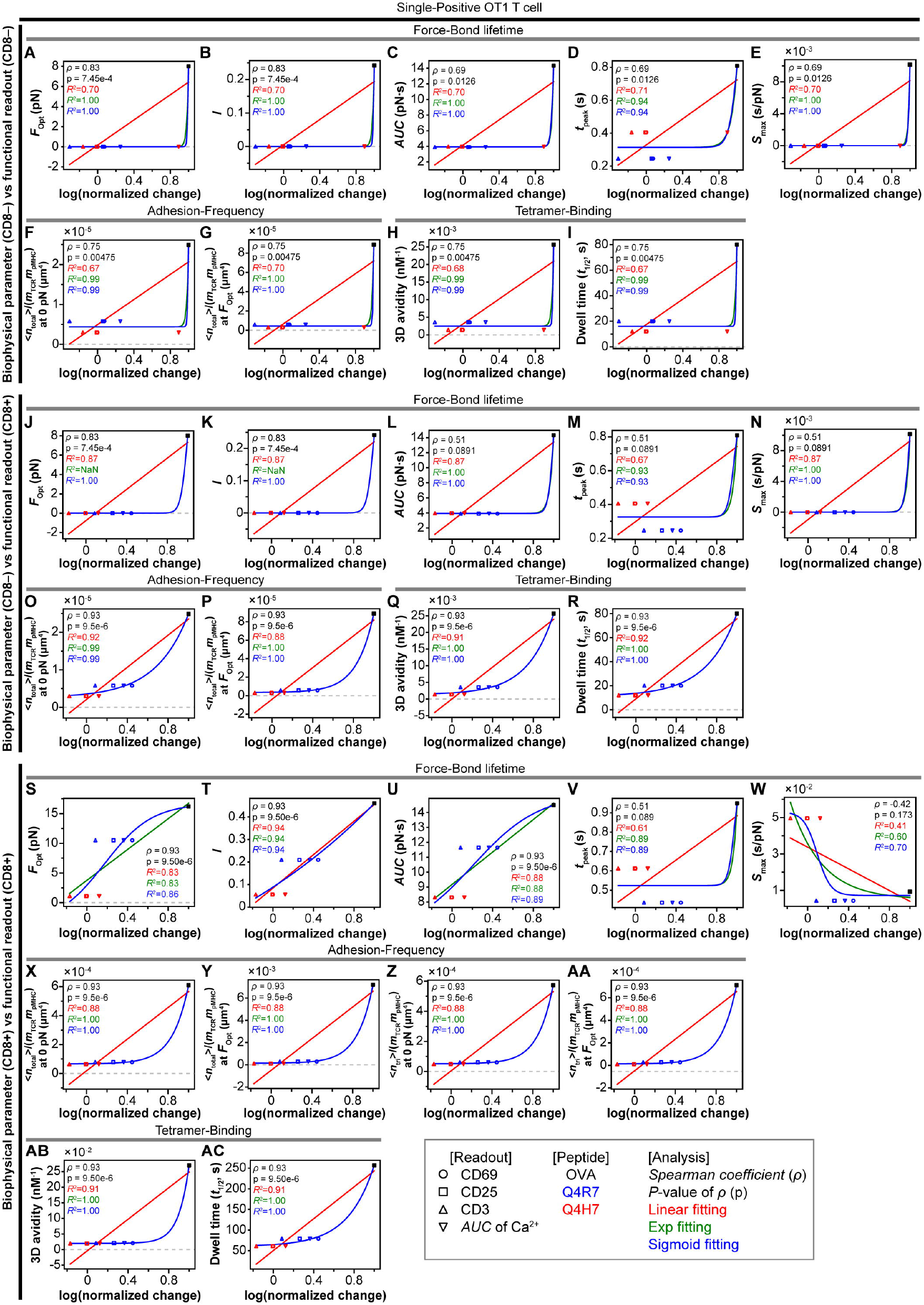

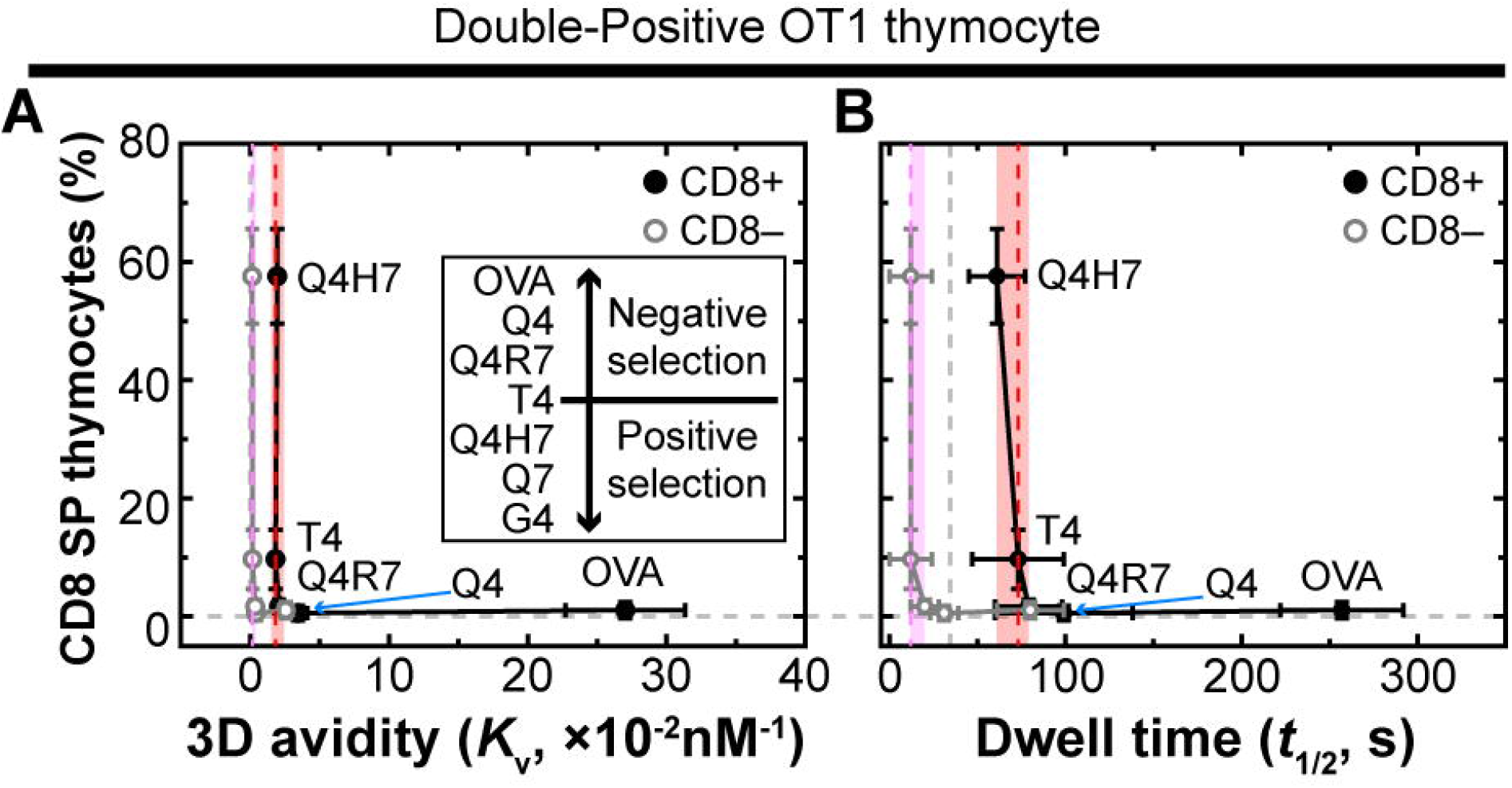

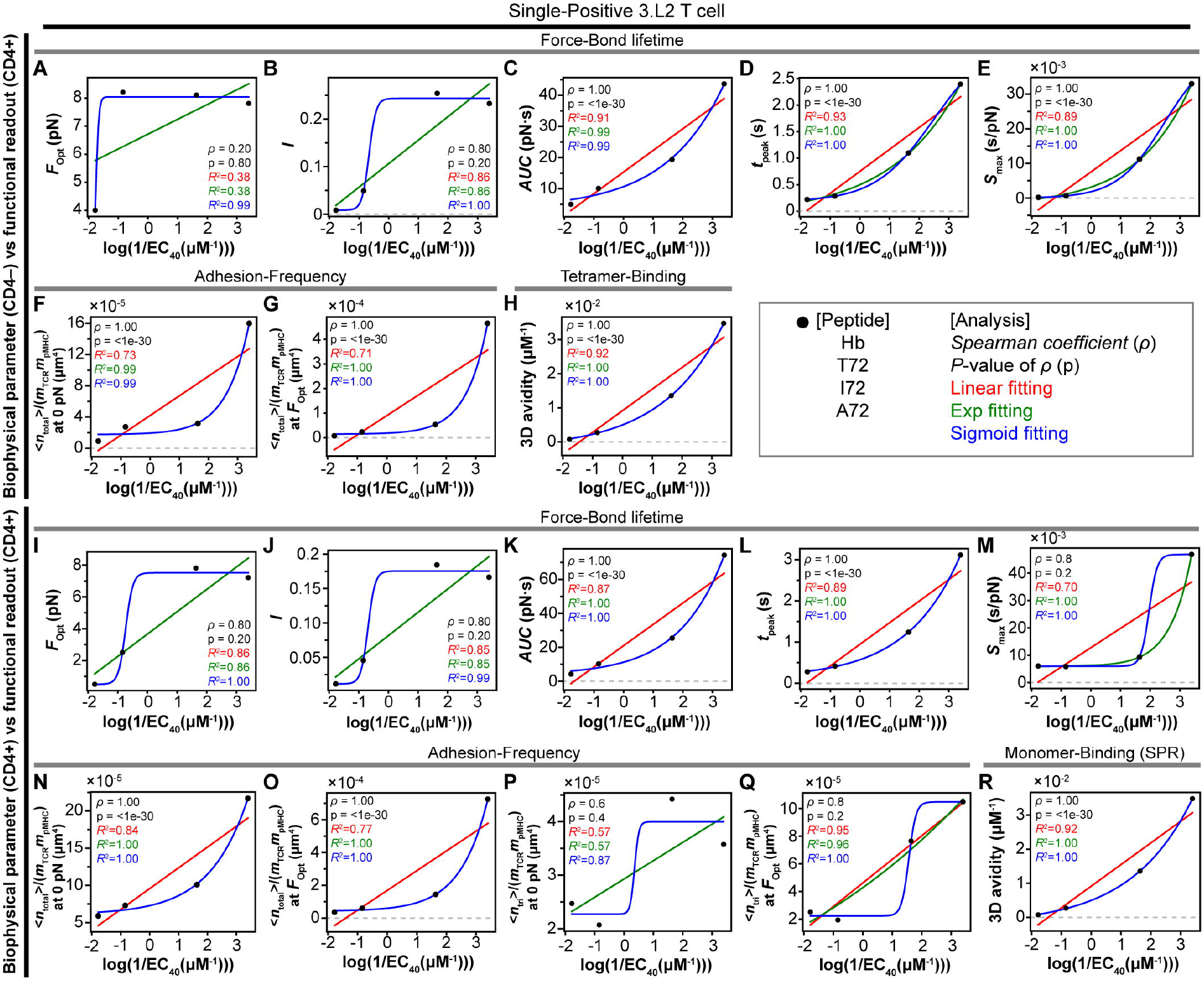

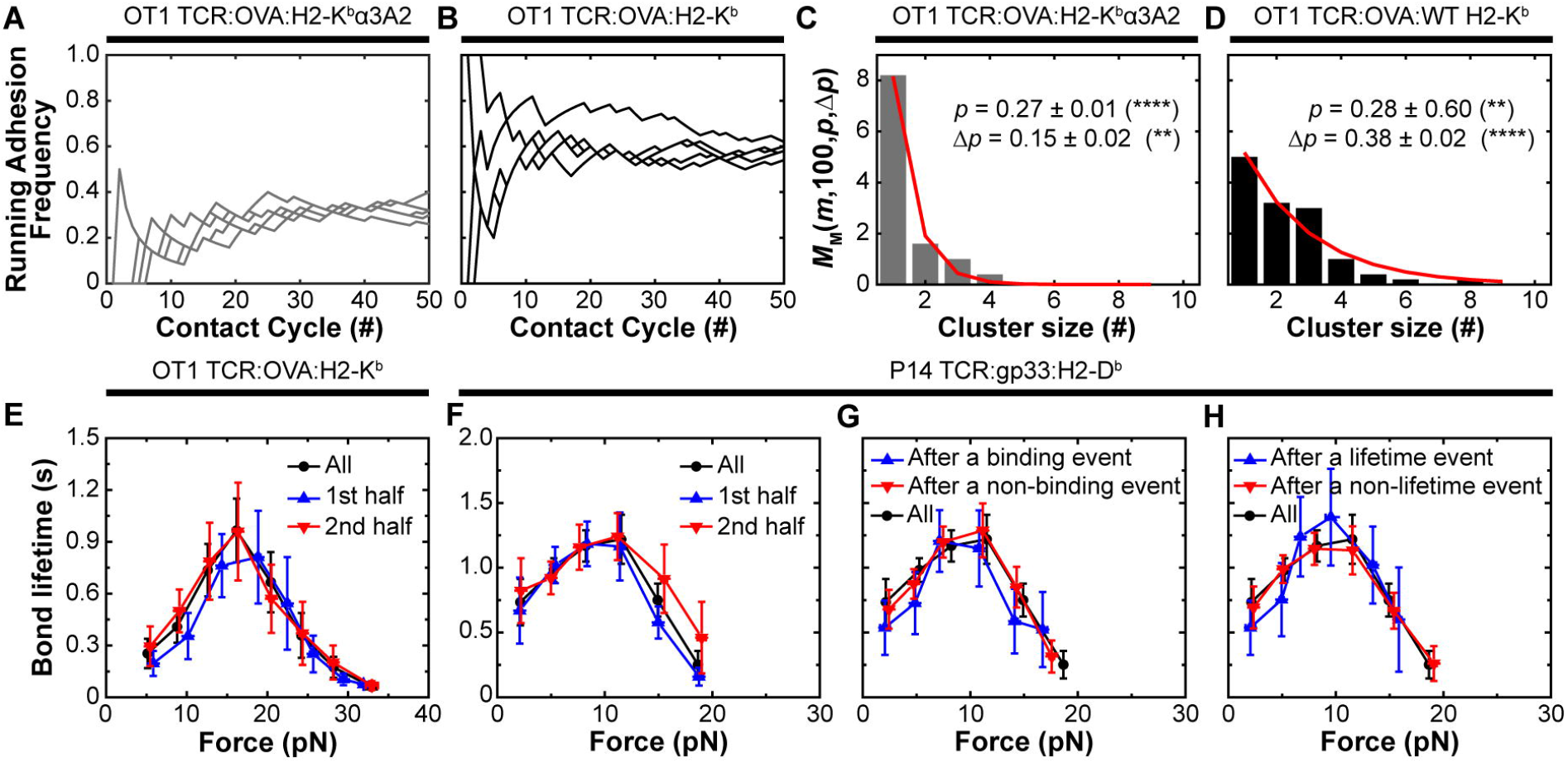

