## Supplementary figures 1 to 9 and Supplementary Table 1 for "Evaluating the effects of CD8/CD4 on T cell function in terms of TCR–pMHC–coreceptor catch and slip bonds"

### *Supplementary Material*

#### 1 Supplementary Figures and Tables

<u>This file includes: Supplementary figures 1 to 9 and Supplementary Table 1</u>

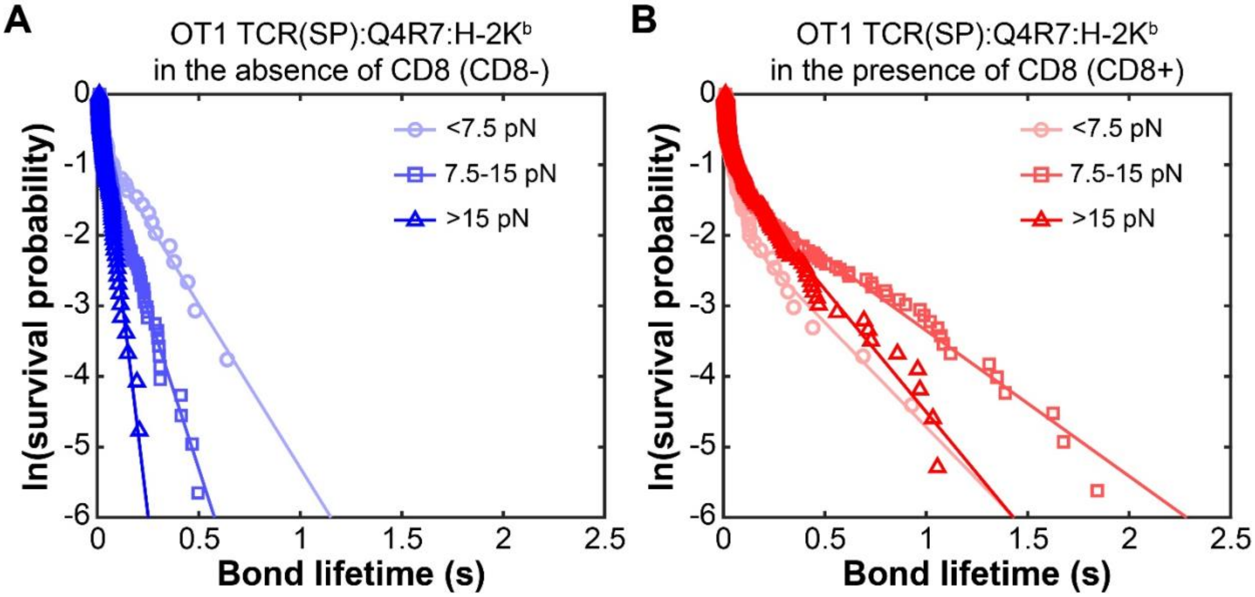


**Supplementary Figure 1. Distributions of lifetimes of bonds of CD8+ OT1 T cell with MT or WT pMHC to prevent or permit CD8 binding at representative forces.**

**(A, B)** *Points:* Semi-log survival probability plots of $\mathbf{ln}$(# of events with a lifetime $> \boldsymbol{t}$) *vs* lifetime *t* of single bonds between OT1 SP naïve T cells and the altered peptide Q4R7 presented by H2-K$^b$α3A2 (A) or H2-K$^b$ (B) at the indicated forces. The curves show pooled ensembles of lifetimes of OT1 TCR–pMHC bonds sorted according to their durations. For each force range, the natural log of the number of events with a lifetime ≥ x-axis value was plotted and fitted by a straight line. The catch and slip bonds can be identified from the changes in decay rate with increasing force in these plots because the steeper the decay, the faster the dissociation.

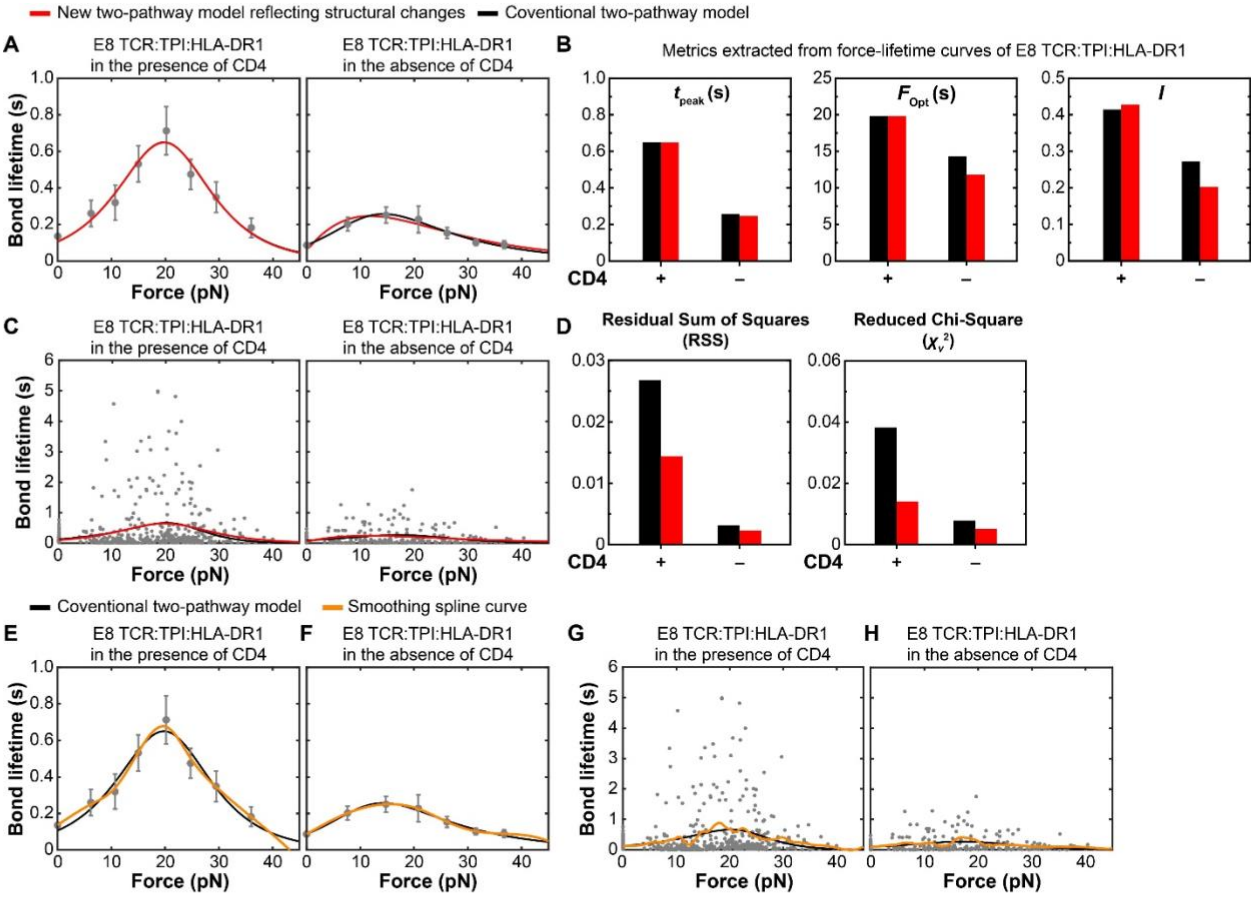


**Supplementary Figure 2. Comparison between the conventional and the structural-based two pathway models.**

**(A)** Fitting of the predicted ($1/k(F)$ curves, *black* = conventional two-pathway model, *red* = structural-based two-pathway model) to the measured (points, Mean ± SEM from n > 20 individual measurements per force bin) lifetime *vs* force curves of the TP1 peptide presented by purified HLA-DR1 interacting with purified E8 TCR in the presence (*left*) and absence (*right*) of CD4. The black curve on the left panel is obscured due to overlapping with the red curve. (**B)** Bar graphs of $t_{\text{peak}}$ (1st panel), $F_{\text{opt}}$ (2nd panel), and $I$ (3rd panel) calculated from the fitted curves shown in (A). **(C)** Fitting of the predicted ($1/k(F)$ curves, *black* = conventional two-pathway model, *red* = structural-based two-pathway model) to individual lifetime *vs* force measurements without binning. **(D**) Comparison of residual sum of squares (RSS, *left*) and reduced Chi-square values ($\chi_v^2$, *right*) obtained from the fitting in (C). Data are from our previous publication (1). **(E, F)** Comparison of fitting by the conventional two-pathway model (*black curves*) and by the splining model (*orange curves*) to the same bin data in A. **(G, H)** Comparison of fitting by the conventional two-pathway model (*black curves*) and by the splining model (*orange curves*) to the same un-binned individual lifetime *vs* force measurements in (C).

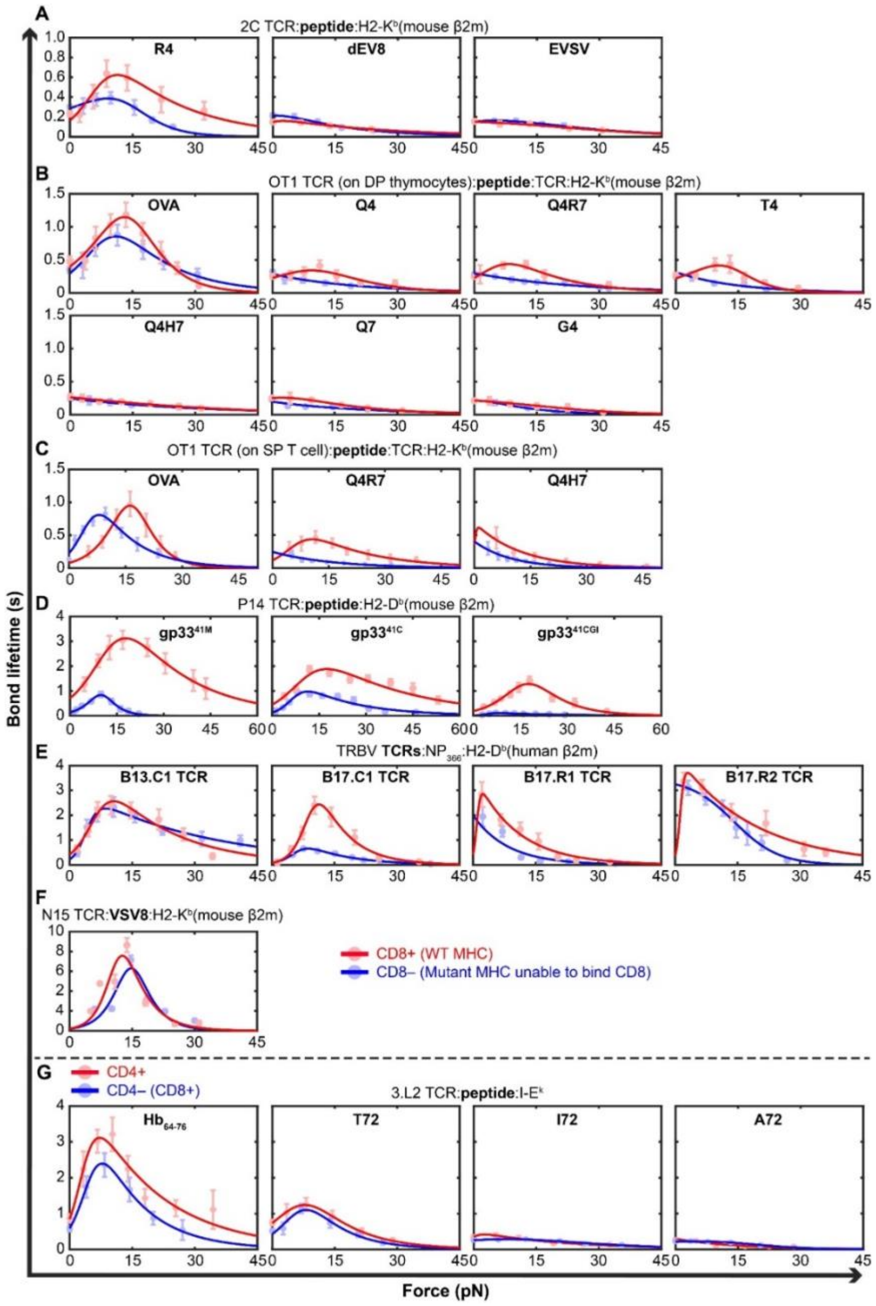


**Supplementary Figure 3. Fitting the conventional two-pathway model to data of lifetime of T cell bonds *vs* force measured in the presence and absence of coreceptor binding.**

**(A-G)** Fitting of theoretical $1/k(F)$ curves predicted by two-pathway model to experimental lifetime (points, Mean ± SEM from at least n > 20 individual measurements per force bin) *vs* force data of 2C (A) (2), DP thymocyte OT1 (B) (2), SP T cell OT1 (C, this study), P14 (D) (3), TRBV (E) (4), and N15 (F) (5) TCRs interacting with the indicated peptides presented by WT or MT H2-$K^b$ or H2-$D^b$ (A-F), and 3.L2 TCR expressed on $CD4^+CD8^-$ or $CD4^-CD8^+$ naïve T cells interacting with the indicated peptides presented by I-$E^k$ (G) (6, 7) to permit (red point and curve) or prevent (blue point and curve) coreceptor from binding. Data are previously published in (2-6). Note that the data in main Fig. 1D are replotted here in (B) (the OVA, T4, and G4 panels) for completeness.

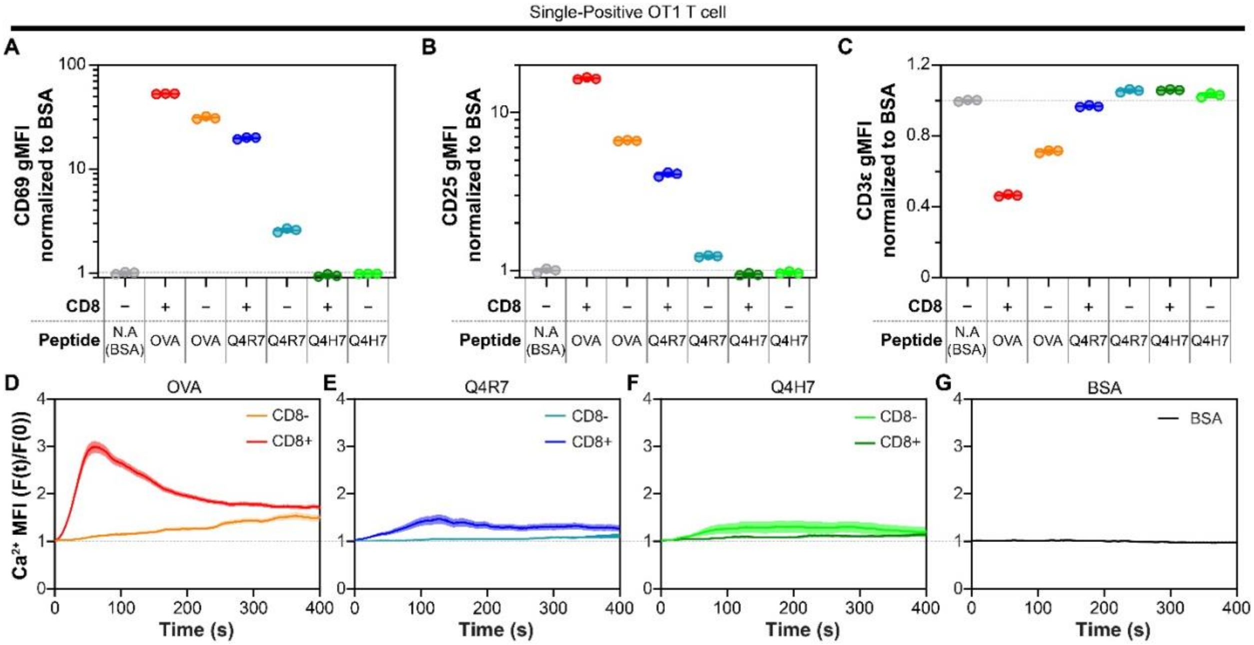


**Supplementary Figure 4. Functional response of SP naïve OT1 T cells to pMHC stimulations.**

**(A-C)** Upon 6 hours stimulation by surfaces coated with 20 µg/ml of BSA (negative control) or 20 µg/ml of OVA, Q4R7, or Q4H7 (indicated) presented by H2-$K^b$ (CD8+) or H2-$K^b$α3A2 (CD8-) at triplicate per condition, the expressions on SP naïve OT1 T cells of CD69 (A), CD25 (B), and CD3 (C) were evaluated by flow cytometry using an fluorescently tagged antibody cocktail (see methods). The geometric mean fluorescence intensity (gMFI) of each experimental sample was normalized by that of BSA. **(D-G)** SP naïve OT1 T cells were loaded with the 5 µM calcium dye (X-Rhod-1) and placed on surfaces coated with 20 µg/ml of OVA (D), Q4R7 (E), or Q4H7 (F) presented by H2-$K^b$α3A2 (CD8-) or H2-$K^b$ (CD8+), or BSA (G), The changes in intracellular calcium signals were measured by live cell imaging using a fluorescence microscope at 0.5s frames per second over 12 mins. The mean fluorescence intensity (MFI) of each cell in each timeframe was normalized by that of its landing frame (set at time = 0 for that cell). Data in (D-G) are presented as mean ± SEM of 59-288 cells.

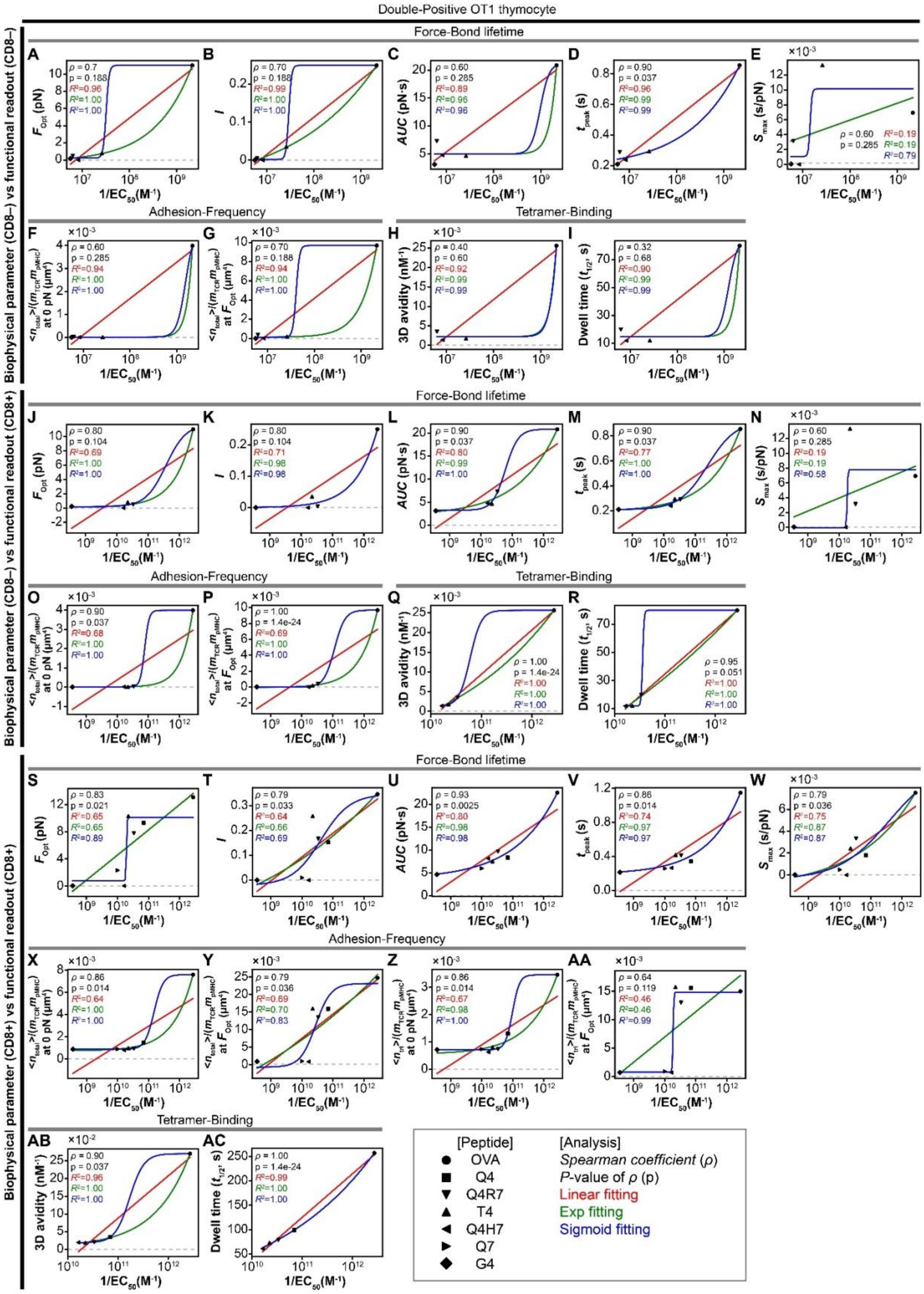

Double-Positive OT1 thymocyte
Force-Bond lifetime
Adhesion-Frequency
Tetramer-Binding
Biophysical parameter (CD8–) vs functional readout (CD8–)
Biophysical parameter (CD8–) vs functional readout (CD8+)
Biophysical parameter (CD8+) vs functional readout (CD8+)
1/EC50(M-1)
[Peptide]
OVA
Q4
Q4R7
T4
Q4H7
Q7
G4
[Analysis]
Spearman coefficient (ρ)
P-value of ρ (p)
Linear fitting
Exp fitting
Sigmoid fitting

**Supplementary Figure 5. Correlative analysis between biophysical metrics of DP OT1 thymocyte interacting pMHC without or with CD8 cooperation and biological activity.**

**(A-AC)** Three combinations of plots were made: 1) $CD8^-$ parameters *vs* $CD8^-$ function (A-I), 2) $CD8^-$ parameters *vs* $CD8^+$ function (J-R), $CD8^+$ parameters *vs* $CD8^+$ function (S-AC). Biophysical metrics of OT1 TCR on DP thymocytes binding to a panel of 5-7 peptides (OVA, Q4R7, T4, Q4H7, G4, in descending order of *x*-axis values) presented by $H2\text{-}K^b\alpha3A2$ or $H2\text{-}K^b$ to prevent or permit CD8 to bind MHC, are plotted *vs* the logarithm of the reciprocal peptide concentration required to stimulate half-maximal CD69 upregulation (1/EC50) and fitted by straight (red), exponential (green), and sigmoidal (blue) curves (different curves sometimes coincide, hence one obscuring the other). Biophysical metrics were measured by BFP force-clamp assay – optimal force $F_{\mathrm{opt}}$ (A J, and S), catch bond intensity $I$ (B, K, and T), area under the curve $AUC$ (C, L, and U), peak bond lifetime $t_{\mathrm{peak}}$ (D, M, and V), and maximum slope $S_{\max}$ (E, N, and W); micropipette adhesion frequency assay – normalized average number of total bonds at zero force $\frac{\langle n\rangle_{\mathrm{tot}}}{(m_{\mathrm{TCR}}m_{\mathrm{pMHC}})}$ (F, O, and X) and at $F_{\mathrm{opt}}$ $\frac{\langle n\rangle_{\mathrm{tot}}(F_{\mathrm{opt}})}{(m_{\mathrm{TCR}}m_{\mathrm{pMHC}})}$ (G, P, and Y), normalized synergy at zero force $\frac{\langle n\rangle_{\mathrm{tri}}}{(m_{\mathrm{TCR}}m_{\mathrm{pMHC}})}$ (Z) and at $F_{\mathrm{opt}}$ $\frac{\langle n\rangle_{\mathrm{tri}}(F_{\mathrm{opt}})}{(m_{\mathrm{TCR}}m_{\mathrm{pMHC}})}$ (AA); tetramer binding – 3D avidity $K_V$ (H, Q, and AB) and dwell time $t_{1/2}$ (I, R, and AC). $R^2$ values for the three curve-fits, shown by matched colors, $P$-values indicating the statistically significant levels of fitting curves, and the Spearman's rank correlation coefficient $\rho$ are shown in each panel to gauge the level of correlation and goodness-of-fit. The CD69 upregulation data are directly from (7). The 3D binding parameters ($K_V$ and $t_{1/2}$) are from (8). The 2D binding parameters are either directly taken from (2) or calculated ($\frac{\langle n\rangle_{\mathrm{tot}}}{(m_{\mathrm{TCR}}m_{\mathrm{pMHC}})}$, $\frac{\langle n\rangle_{\mathrm{tri}}}{(m_{\mathrm{TCR}}m_{\mathrm{pMHC}})}$, $F_{\mathrm{opt}}$, $I$, $AUC$, $t_{\mathrm{peak}}$, and $S_{\max}$ by fitting the data from (2) using the two-pathway model (cf. Supplementary Figure 3). Error bars are either directly from experimental data or calculated from experimental data from (2, 7, 8).

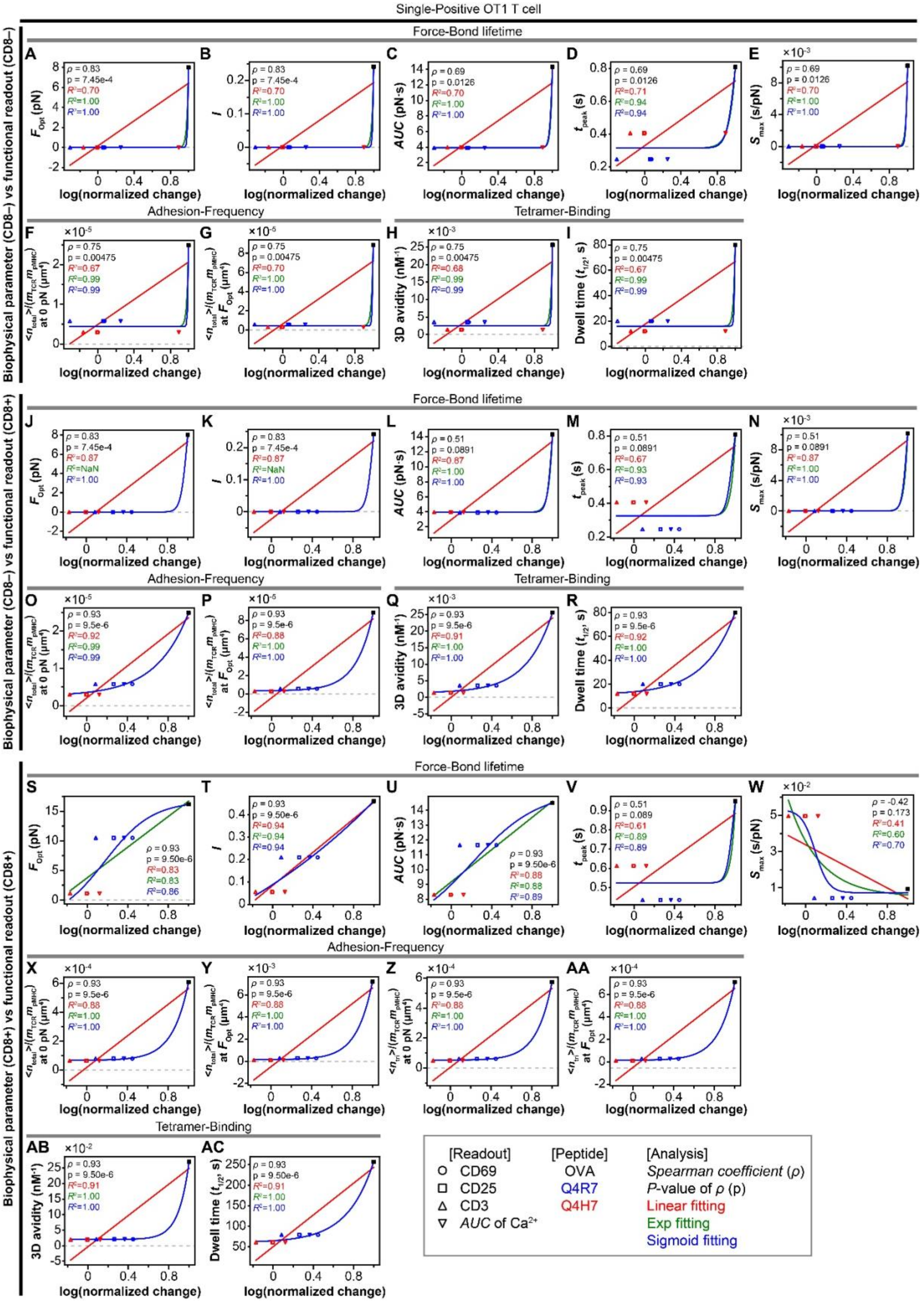

Single-Positive OT1 T cell
Biophysical parameter (CD8–) vs functional readout (CD8–)
Force-Bond lifetime
Adhesion-Frequency
Tetramer-Binding
Biophysical parameter (CD8–) vs functional readout (CD8+)
Force-Bond lifetime
Adhesion-Frequency
Tetramer-Binding
Biophysical parameter (CD8+) vs functional readout (CD8+)
Force-Bond lifetime
Adhesion-Frequency
Tetramer-Binding
log(normalized change)
[Readout]
CD69
CD25
CD3
AUC of Ca2+
[Peptide]
OVA
Q4R7
Q4H7
[Analysis]
Spearman coefficient (ρ)
P-value of ρ (p)
Linear fitting
Exp fitting
Sigmoid fitting

**Supplementary Figure 6. Correlating biophysical metrics of SP OT1 naïve T cells interaction with pMHC without or with CD8 cooperation and in the absence or presence of force with biological activity.**

**(A-AC)** Three combinations of plots were made: 1) $CD8^-$ parameters *vs* $CD8^-$ function (A-I), 2) $CD8^-$ parameters *vs* $CD8^+$ function (J-R), $CD8^+$ parameters *vs* $CD8^+$ function (S-AC). Biophysical metrics of OT1 TCR on SP naïve T cells binding to a panel of 3 peptides (OVA = black, Q4R7 = blue, Q4H7 = red) presented by H2-$K^b\alpha$3A2 or H2-$K^b$ to prevent or permit CD8 bind MHC, are plotted *vs* the logarithm of normalized changes in expression of CD69 (circle), CD25 (square), CD3 (triangle), and intracellular calcium concentration (inverted triangle), and fitted by straight (red), exponential (green), and sigmoidal (blue) curves (different curves sometimes coincide, hence one obscuring the other). Biophysical metrics were measured by BFP force-clamp assay – optimal force $F_{\mathrm{opt}}$ (A J, and S), catch bond intensity $I$ (B, K, and T), area under the curve $AUC$ (C, L, and U), peak bond lifetime $t_{\mathrm{peak}}$ (D, M, and V), and maximum slope $S_{\mathrm{min}}$ (E, N, and W); micropipette adhesion frequency assay – normalized average number of bonds at zero force $\frac{\langle n\rangle_{\mathrm{tot}}}{(m_{\mathrm{TCR}}m_{\mathrm{pMHC}})}$ (F, O, and X) and at $F_{\mathrm{opt}}$ $\frac{\langle n\rangle_{\mathrm{tot}}(F_{\mathrm{opt}})}{(m_{\mathrm{TCR}}m_{\mathrm{pMHC}})}$ (G, P, and Y), normalized synergy at zero force $\frac{\langle n\rangle_{\mathrm{tri}}}{(m_{\mathrm{TCR}}m_{\mathrm{pMHC}})}$ (Z) and at $F_{\mathrm{opt}}$ $\frac{\langle n\rangle_{\mathrm{tri}}(F_{\mathrm{opt}})}{(m_{\mathrm{TCR}}m_{\mathrm{pMHC}})}$ (AA); tetramer binding – 3D avidity $K_V$ (H, Q, and AB) and dwell time $t_{1/2}$ (I, R, and AC). $R^2$ values for the three curve-fits, shown by matched colors, $p$-values indicating the statistically significant levels of fitting curves, and the Spearman's rank correlation coefficient $\rho$ are shown in each panel to gauge the level of correlation and goodness-of-fit. The T cell functional responses are normalized from the data shown in Supplementary Figure 4. The 3D binding parameters ($K_V$ and $t_{1/2}$) are from (8). The five catch bond metrics $F_{\mathrm{opt}}$, $I$, $AUC$, $t_{\mathrm{peak}}$, and $S_{\mathrm{max}}$ are determined by first fitting the force-dependent lifetime data shown in Supplementary Figure 3C using the two-pathway model and then calculating values based on their definition shown in Figure 1B.

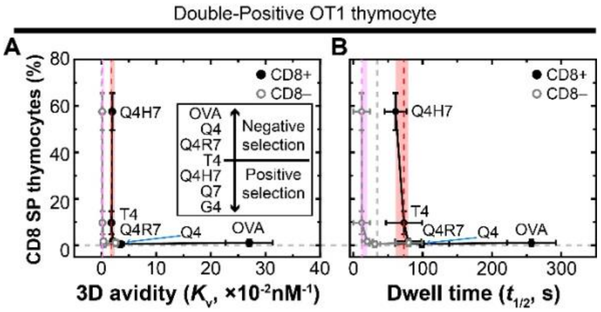


**Supplementary Figure 7. CD8 impacts TCR specificity for two ligands evaluated by 3D measures.**

**(A, B)** The percentage of $CD8^+$ SP thymocytes from FTOC assay are plotted *vs* 3D avidity $K_V$ (A) or dwell time $t_{1/2}$ (B) of OT1 TCR interacting with the same panel of 7 peptides presented by H2-$K^b$ ($CD8^+$, closed black circles connected by black line segments) or H2-$K^b$α3A2 ($CD8^-$, open gray circles connected by gray line segments) to permit or prevent CD8 from binding to MHC. Vertical dashed lines identify the parameters of the threshold peptide (T4) and the vertical stripes mark the parameter ranges across the threshold from the strongest positive selection peptide (Q4H7) to the weakest negative selection peptide (Q4R7) using different colors to indicate measurements made when CD8 was prevented (purple) or permitted (red) to bind MHC. All data of 3D avidity, dwell time, and FTOC assays are from the literature (9).

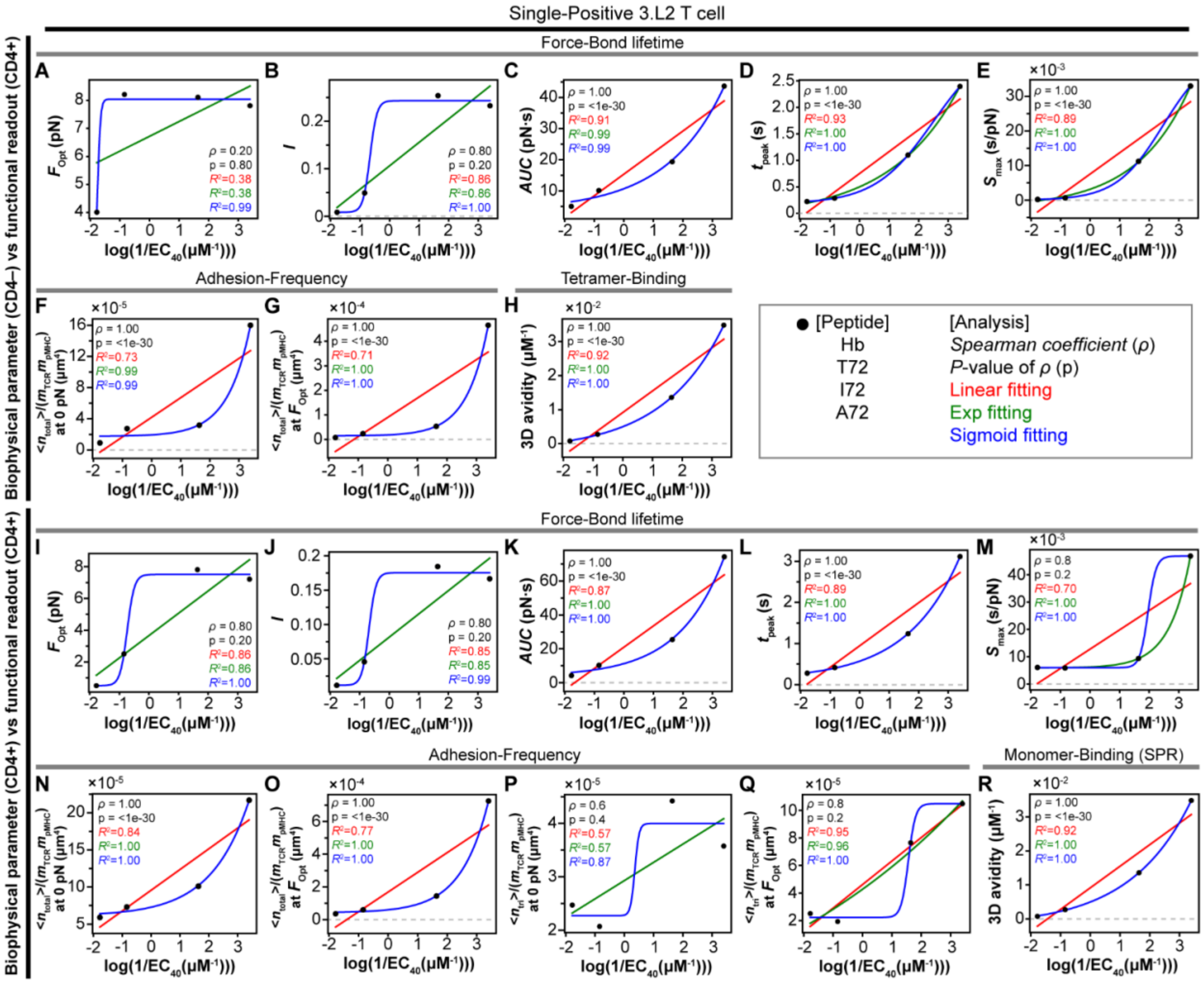


**Supplementary Figure 8. Correlating biophysical metrics of 3.L2 TCR T cells interaction with pMHC without or with CD4 cooperation and in the absence or presence of force with biological activity.**

**(A-Q)**, Biophysical metrics measured using T cells from 3.L2 TCR transgenic mice respectively expressing CD4⁻CD8⁺ to prevent (CD4-, A-H) or CD4⁺CD8⁻ to permit (CD4+, I-R) CD4 to bind I-E$^k$ that presents WT peptide Hb$_{64\text{-}78}$ and its MTs T72, I72, and A72, by micropipette adhesion frequency assay (F, G, N, O, P, Q) and BFP force-clamp assay (A-E and I-M) – effective 2D affinity at zero-force $A_cK_a$ (F, N), effective 2D affinity at optimal force $A_cK_a(F_{\text{opt}}) = A_cK_a \times k_{\text{off}} \times t_{\text{peak}}$ (G, O), normalized synergy at zero force $\frac{\langle n\rangle_{\text{tri}}}{(m_{\text{TCR}}m_{\text{pMHC}})}$ (P) and at $F_{\text{opt}}$ $\frac{\langle n\rangle_{\text{tri}}(\text{at } F_{\text{opt}})}{(m_{\text{TCR}}m_{\text{pMHC}})}$ (Q), optimal force $F_{\text{opt}}$ (A, I), catch bond intensity $I$ (B, J), area under the curve $AUC$ (C, K), peak bond lifetime $t_{\text{peak}}$ (D, L), and maximum slope $S_{\max}$ (E, M) – are plotted vs the logarithm of the reciprocal peptide concentration required to stimulate 40% maximal IL2 (1/EC40) and fitted by straight (red), exponential (green) and sigmoidal (blue) curves (different curves sometimes coincide, hence one obscuring the other). $R^2$ values for the three curve-fits, shown by matched colors, $P$-values indicating the statistically significant levels of fitting curves, and the Spearman's rank correlation coefficient $\rho$ are shown in each panel to gauge the level of correlation and goodness-of-fit. The data in the plots were from re-analysis of the original data published in (1, 6). **(R)** The same correlative analysis was performed using the 3D avidity as the ***y***-axis variable from (7).

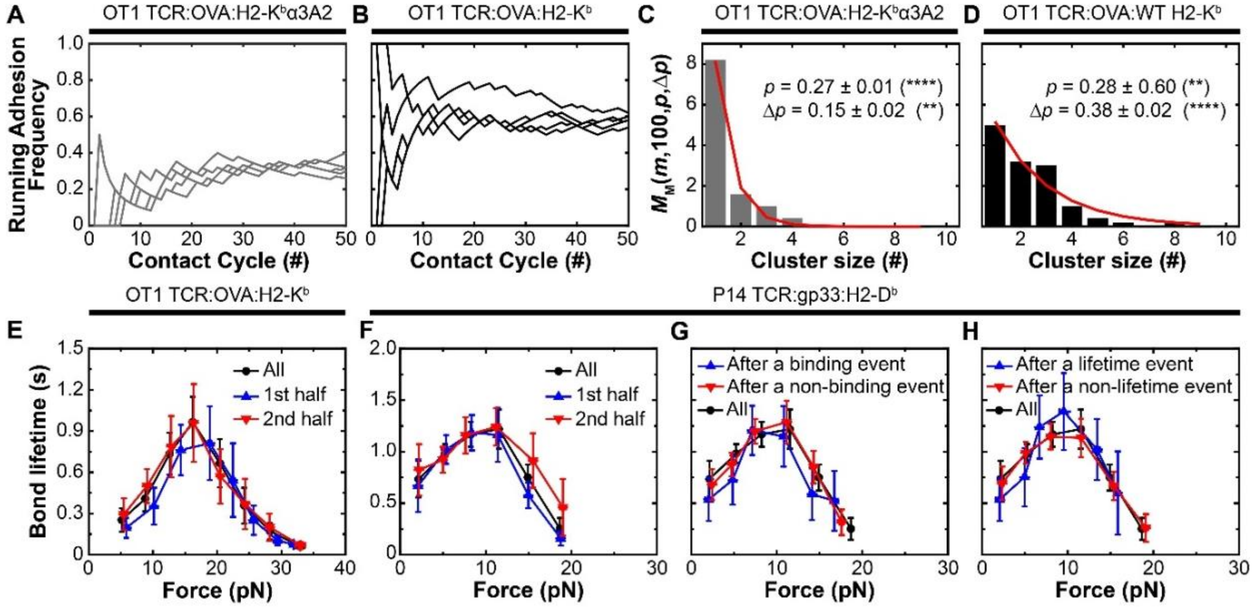


**Supplementary Figure 9. Analysis of molecular memory in the adhesion frequency and force-dependent bond lifetimes.**

**(A, B)** Representative running adhesion frequency *vs* cycle number of repeated contacts between individual naïve OT1 T cells and red blood cells coated with OVA peptide presented by MT H2-$K^b$α3A2 (A) or WT H2-$K^b$ (B). (**C, D**) Adhesion cluster size distributions (bar) and model fit (curve) calculated from the above running adhesion frequency data obtained in the absence (C) and presence (D) of CD8 contribution using the method described in (10, 11). $^{**}P<0.01$, $^{****}P<0.0001$ indicating the statistically significant levels of fitting curves. **(E, F)** Mean ± SEM of single bond lifetimes of OT1 naïve T cells interacting with $OVA_{257\text{-}264}$:H2-$K^b$ (E) or P14 naïve T cells interacting with $gp33^{41M}$:H2-$D^b$ (F) measured from the first half (blue triangle), last half (red inverted triangle), and all (black circle) lifetime events in the time series generated by repetitively testing a single cell. **(G, H)** Mean ± SEM of lifetimes of single P14 TCR–$gp33^{41M}$:H2-$D^b$ bonds measured after a binding (G) or lifetime (H) event (blue triangle), or after a non-binding (G) or non-lifetime (H) event (red inverted triangle), and all events (black circle). Data are pooled from 10 (E) and 30 (F-H) cells with 5-25 measurements per cell.

| Super agonist ~ Agonist | | | Weak agonist | | | Antagonist | | |
|---|---|---|---|---|---|---|---|---|
| TCR | peptide | MHC I* | TCR | peptide | MHC I* | TCR | peptide | MHC I* |
| N15 | VSV8 | H-2K$^b$ | OT1 | Q4 | H-2K$^b$ | OT1 | G4 | H-2K$^b$ |
| P14 | gp33$^{41M}$ | H-2D$^b$ | OT1 | Q4R7 | H-2K$^b$ | 2C | EVSV | H-2K$^b$ |
| P14 | gp33$^{41C}$ | H-2D$^b$ | OT1 | T4 | H-2K$^b$ | P14 | gp33$^{41CGI}$ | H-2D$^b$ |
| OT1 | OVA | H-2K$^b$ | OT1 | Q4H7 | H-2K$^b$ | B17.R1$^\dagger$ | NP366 | H-2D$^b$ |
| 2C | R4 | H-2K$^b$ | OT1 | Q7 | H-2K$^b$ | B17.R2$^\dagger$ | NP366 | H-2D$^b$ |
| 2C | Dev8 | H-2K$^b$ | P14 | gp33$^{41C}$ | H-2D$^b$ | | | |
| B13.C1 | NP366 | H-2D$^b$ | | | | | | |
| B17.C1 | NP366 | H-2D$^b$ | | | | | | |

**Supplementary Table 1. Summary of TCRs and pMHC-Is and their classification.**

*MHCs include WT and mutant (α3A2) to abolish CD8 binding.

†Reverse-docking topology of TCRs are considered as antagonist albeit the peptide (NP366) is known as agonist for canonical-docking topology of TCR binding to MHC.
